# Translational contributions to tissue-specificity in rhythmic and constitutive gene expression

**DOI:** 10.1101/060368

**Authors:** Violeta Castelo-Szekely, Alaaddin Bulak Arpat, Peggy Janich, David Gatfield

## Abstract

**BACKGROUND:** The daily gene expression oscillations that underlie mammalian circadian rhythms show striking tissue differences and involve post-transcriptional regulation. Both aspects remain poorly understood. We have used ribosome profiling to explore the contribution of translation efficiency to temporal gene expression in kidney, and contrasted our findings with liver data available from the same mice.

**RESULTS:** Rhythmic translation of constantly abundant mRNAs affected largely non-overlapping transcript sets with distinct phase clustering in the two organs. Moreover, tissue differences in translation efficiency modulated the timing and amount of protein biosynthesis from rhythmic mRNAs, consistent with organ-specificity in clock output gene repertoires and rhythmicity parameters. Our comprehensive datasets provided insights into translational control beyond temporal regulation. Between tissues, many transcripts showed differences in translation efficiency, which were, however, of markedly smaller scale than mRNA abundance differences. Tissue-specific changes in translation efficiency were associated with specific transcript features and, intriguingly, globally counteracted and compensated transcript abundance variations, leading to higher similarity at the level of protein biosynthesis between both tissues.

**CONCLUSIONS:** We show that tissue-specificity in rhythmic gene expression extends to the translatome and contributes to define the identities, the phases and the expression levels of rhythmic protein biosynthesis. Moreover, translational compensation of transcript abundance divergence leads to overall higher similarity at the level of protein production across organs. The unique resources provided through our study will serve to address fundamental questions of post-transcriptional control and differential gene expression *in vivo*.

## Background

Circadian clocks serve organisms to synchronize behaviour, physiology and gene expression according to time of day. The mammalian circadian system consists of a master clock in the brain’s suprachiasmatic nuclei (SCN) that receives photic inputs from the retina and synchronizes peripheral clocks present in most cells throughout the body. The molecular timekeeping mechanism – the core clock – consists of a network of transcriptional activators and repressors interacting in negative feedback loops (reviewed in [1, 2]). In the core loop, the heterodimeric transcription factor ARNTL:CLOCK (also known as BMAL1:CLOCK) drives the expression of its own repressors, encoded by the *Period (Per1, Per2, Per3)* and *Cryptochrome (Cry1, Cry2)* genes – a configuration also known as the positive and negative limbs of the oscillator. Additional feedback – in particular an interconnecting limb involving nuclear receptors of the REV-ERB (encoded by genes *Nr1d1, Nr1d2*) and ROR *(Rora, Rorb, Rorc)* family – intersects with the core loop, and numerous post-translational modifications of clock proteins further add to the complexity of the circuitry. The final outcome is a set of robustly cycling transcriptional activities peaking at different phases around the day that drive the rhythmic expression of hundreds to thousands of other genes, termed the clock output or clock-controlled genes (CCGs). It is noteworthy that, despite the probably (near-)identical molecular makeup of the core clock across cell types, CCGs show considerable tissue-specificity [3]. The co-regulation by core clock and tissue-specific (non-rhythmic) transcription factors may engender such cell type-specific rhythmic expression patterns, as shown to occur in *Drosophila* [4]. Overall, however, the origins of tissue-specificity in rhythmic gene output (and even in certain core clock parameters [5]) are poorly understood. Mechanisms that act at the post-transcriptional level and that impact daily mRNA and protein accumulation kinetics are plausible players in the generation of cell type differences as well.

Rhythmic gene expression has been mainly investigated at the transcriptome level i.e., using mRNA abundances as a primary readout. However, comparison of mRNA levels with datasets of genome-wide transcriptional activity and of protein abundances that have become available recently, has suggested that a surprisingly large fraction of gene expression oscillations may have post-transcriptional origins (reviewed in [6]). The many cases of protein rhythms that are independent of an underlying oscillating transcript (initially reported in a low-throughput mass-spectrometric study from mouse liver already 10 years ago [7] and recently confirmed at a comprehensive scale [8, 9]) point to important roles for translation, protein degradation and protein secretion in shaping time of day-dependent proteomes. We [10] and others [11] have recently used ribosome profiling, a genome-wide method that assesses translation efficiency through the deep sequencing of ribosome-protected mRNA fragments, to chart the contribution of translational control to daily protein biosynthesis in mouse liver. One conclusion that emerged from the identified cases of translationally generated oscillations was that circadian clock activity and feeding rhythms both contribute to regulating rhythmic gene expression outputs [10, 11]. Notably, the most abundant group of transcripts subject to rhythmic translation, i.e. mRNAs encoding ribosomal proteins and other components of the translation machinery that all contain 5′-terminal oligopyrimidine tract (5′-TOP) sequences regulated by the mammalian target of rapamycin (mTOR) [12], appear to be under the dominant control of feeding [11].

We have now performed ribosome profiling using a second organ from the same cohort of animals, the kidney, which is an emerging circadian model organ with distinct rhythmic functions [13]. By contrasting kidney and liver datasets we comprehensively assessed commonalities and differences in their translatomes, and we evaluated in how far the regulation of translation efficiency contributed to tissue specificity in rhythmic and constitutive protein biosynthesis.

## Results

### Around-the-clock ribosome profiling datasets from two organs

For our recent study of the liver translatome around-the-clock [10] we had used ribosome profiling [14] (RPF-seq) on a time series of organs collected from mice sacrificed every 2 hours over the 24-hour day (12 timepoints in duplicate; Figure 1A). To generate a complementary dataset from a second organ we chose the kidneys from the same cohort of animals. Liver and kidney express thousands of genes in common [3, 15], thus providing a particularly suitable setting for a cross-organ comparison of gene expression.

**Fig. 1.**
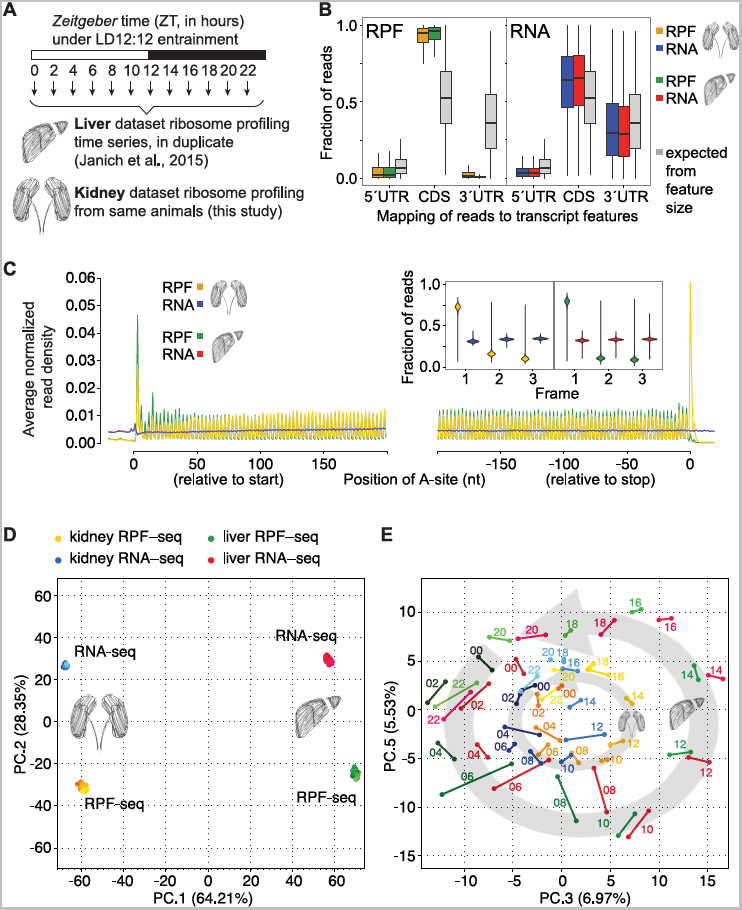
Ribosome profiling around-the-clock in mouse liver and kidney. **A** Overview of the experimental design. Livers and kidneys for ribosome profiling were collected every 2 hours for two daily cycles. Each timepoint sample was a pool of organs from two animals. Mice were kept under 12 hour:12 hour light-dark conditions, with *Zeitgeber* times ZT00 corresponding to lights-on and ZT12 to lights-off. **B** Read distribution to transcript features. RPF-seq (left; kidney in orange, liver in green) and RNA-seq (right; blue and red for kidney and liver, respectively) compared to a distribution expected from the relative feature sizes (grey; the distributions based on feature sizes were highly similar for both organs, thus only that for kidney is shown). Note that RPF-seq footprints were enriched on the CDS and depleted from UTRs, whereas RNA-seq reads distributed more homogeneously along transcripts, according to feature size. Of note, the higher level of 3′ UTR footprints in kidney resulted mainly from differences in the efficiency with which stop codon footprints were captured, as described in panel C. **C** Predicted position of the ribosome’s aminoacyl tRNA-site (A-site) of reads relative to the CDS start and stop codons. Read density at each position was averaged across single protein coding isoform genes (i.e., genes with one main expressed transcript isoform) that had an average RPF RPKM >5, a CDS >400 nt in length and were expressed in both organs (N=3037 genes). This analysis revealed the trinucleotide periodicity of RPF-seq (but not RNA-seq) reads in both organs. Inset: Frame analysis of CDS reads showed preference for the annotated reading frame (frame 1, the same frame as the start codon) in RPF but not in RNA reads. Violin plots extend to the range of the data (N=3694 genes for liver, N=4602 genes for kidney). A separate analysis of the higher level of stop codon footprints in kidney, that also led to the differences in 3′ UTR reads in B, can be found in Figure S2A-B. **D** Principal component (PC) analysis of kidney and liver RPF-seq and RNA-seq datasets, using the 4000 most variable genes. The first two components reflected the variability coming from organ (PC1, 64.21%) and from RPF/RNA origin of datasets (PC2, 28.35%). **E** PC3 vs. PC5 (together 12.5% of variation) resolved the factor time within each dataset, leading to a representation that resembled the face of a clock. Each dot represents one sample, timepoint replicates are joined by a line and timepoints within each dataset are sequentially coloured. The circular arrangement was larger for liver than kidney, suggesting a higher contribution of hepatic rhythmic genes to overall variability. Figure S4 shows the scree plot for the ten first components.

Applying the same experimental and computational methods as for liver RPF-seq [10, 16], we obtained comparable high-quality data for kidney (see Supplementary Table S1 and Figure S1A-C for details on sequencing and mapping outcomes). Briefly, ribosome footprints from both organs showed similar enrichment for protein coding sequences (CDS) of mRNAs and depletion of untranslated regions (UTRs) (Figure 1B). Like the footprints from liver, also those from kidney exhibited excellent reading frame preference, which allowed resolving the 3-nt periodicity of coding sequences transcriptome-wide (Figures 1C and S2A-B). Moreover, the high correlation coefficients seen across replicates of the kidney time series for both RNA- and RPF-seq data indicated excellent biological and technical reproducibility (Figure S3A-B). We also used a recently developed tool, termed Ribo-seq Unit Step Transformation (RUST) [17], to confirm high technical similarity of datasets between organs (Figure S2C-D). Finally, principal component (PC) analysis on all available datasets (96 libraries, i.e. RPF-seq and RNA-seq from 2 organs, 12 timepoints, in duplicate) segregated the data according to the main experimental and biological covariates. PC1 (explaining 64.2% of variation) thus separated libraries according to organ, indicating that tissue origin represented the major source of divergence, followed by PC2 (28.4%) that separated RNA-seq (mRNA abundance) and RPF-seq (footprints/translation) (Figure 1D). The cyclic nature of the data was resolved in the representation PC3 vs. PC5 (together 12.5%), in which timepoints assembled to a near-perfect clock (Figure 1E). The larger circular arrangement of the liver vs. kidney time series suggested that rhythmic gene expression from liver contributed more strongly to overall variation than did kidney rhythms. This observation is in line with the notion that there are more, and higher amplitude rhythms in liver than in kidney [3]. Taken together, we concluded that the kidney data were of similarly high quality as our previous liver datasets [10]. Together, they would be suitable for comparative analyses of time of day-dependent and constitutive translation across two tissues.

### Cross-organ differences in translation efficiency are widespread, of moderate scale, and partially compensate RNA abundance differences

To what extent do differences in translation efficiency contribute to different gene expression outputs across organs? We addressed this question using the set of 10289 genes whose expression was detectable in kidney and in liver at both RPF and RNA level (Figure 2A). From the ratio of CDS-mapping normalised read counts for RPF-seq relative to RNA-seq we first calculated relative translation efficiencies (TEs) per transcript and for each organ. TEs were overall rather similar between tissues, with 95% of genes falling into a less than 3-fold range for the kidney/liver TE ratio, as compared to a greater than 100-fold range for the transcript abundance ratio (Figure 2B). This observation was coherent with the considerably broader spread of mRNA abundances vs. TEs across genes within each organ (greater than 500-fold vs. just over 10-fold, respectively; Figure S5A-B) and is in line with a dominant role for the regulation of mRNA levels (i.e., transcription and mRNA decay) in controlling quantitative differences in gene output.

**Fig.2.**
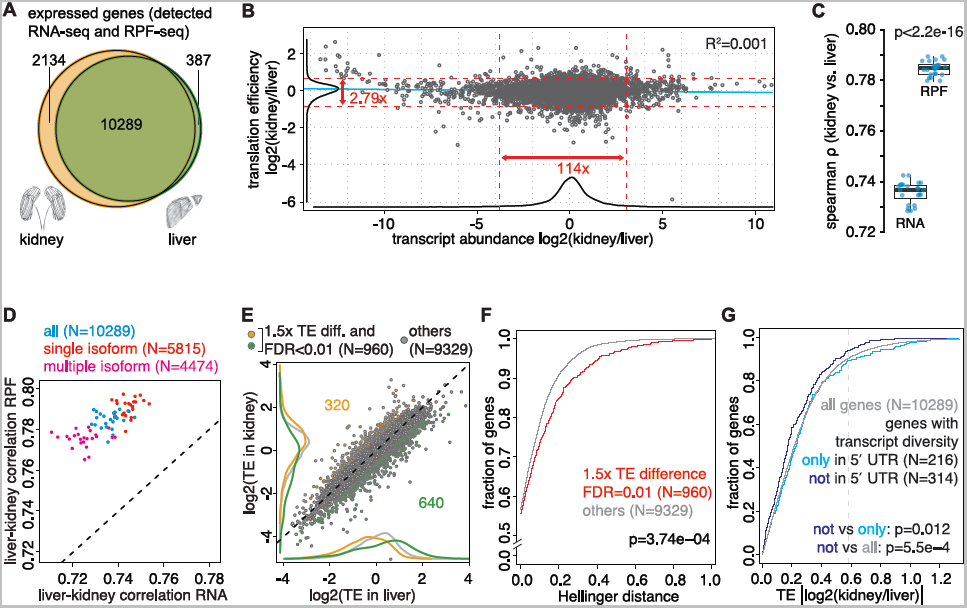
Cross-organ differences in translation efficiency partially compensate RNA abundance differences and show association with transcript features. **A** Venn diagram showing the gene expression overlap (i.e. genes detected at both RPF and RNA level) between kidney (yellow, N=12423 genes) and liver (green, N=10676 genes). Same cut-offs on RPKM (reads per kilobase of transcript per million mapped reads) were used for both organs. **B** Scatterplot of kidney-to-liver ratio of mRNA abundance vs. translation efficiency (TE) for all expressed genes (N=10289), averaged over all timepoints. Corresponding density curves are plotted on the margins. Dashed red lines represent the 2.5 and 97.5 percentiles of each variable, and the corresponding fold-change is indicated. Linear regression line is depicted in blue (R^2^=0.0009, p=0.0009). While 95% of genes spanned a 114-fold range in mRNA abundance differences across organs, the same number of genes changed less than 3-fold in TE, underlining that transcript abundance was the main contributor to divergent gene expression. **C** Inter-organ Spearman correlation for RNA-seq and RPF-seq samples. Each dot represents the correlation coefficient between kidney and liver for a timepoint and replicate sample. Note that RPF-seq samples consistently correlated significantly better than RNA-seq samples did (p<2.2e-16, N=24, paired t-test of Fisher-transformed correlation coefficients). **D** Scatterplot of inter-organ RNA vs. RPF correlation coefficients for each sample separately calculated from all (blue, N=10289), from single isoform (red, N=5815), and from multiple isoform (pink, N=4474) genes. Consistently better RPF correlation was evident in all cases. **E** Relative TE in liver vs. kidney (data centred and averaged over all timepoints for all expressed genes, N=10289) showed an overall strong inter-organ correlation. Differential TE – defined as having FDR-corrected p<0.01 (Wilcoxon signed rank test on TE) and >1.5 difference in TE across organs – was apparent for ca. 9% of genes (yellow and green show cases where TE is higher in kidney and liver, respectively, N=960). **F** Cumulative distribution of Hellinger distances for genes showing differential TE (red, N=960), or not (grey, N=9329), as detected in (E). Hellinger distance was used as a quantitative measure for relative transcript isoform diversity across organs, as described in Results and Methods. The analysis shows that divergent TE correlated with larger diversity in transcript isoform expression (D=0.0702, p=3.74e-04, two-sample Kolmogorov-Smirnov [KS] test). **G** Cumulative distribution of the kidney-to-liver TE ratio for genes whose transcript diversity originated exclusively from the 5′ UTR (identical CDS and 3' UTR, light blue, N=216; these genes show more TE differences across organs), and genes whose transcripts had identical 5′ UTR (and divergent CDS and/or 3′ UTR, purple, N=314; these genes show less TE differences across organs). The vertical dashed grey line marks the 1.5-fold difference used to define differential TE (as in (E)). These results suggested that tissue specificity in TE was partially achieved by expressing transcript isoforms that differed in their 5′ UTRs (note the significant shift towards smaller TE differences for genes with identical 5' UTRs). See also Figure S9.

Intuitively, we had expected that RNA levels that were widely dissimilar between kidney and liver and subsequently further modulated by organ-specific TEs, would probably give rise to even greater cross-organ divergence at the RPF level. Intriguingly, however, the global correlation between kidney and liver was better for footprint abundances than for transcript abundances (Spearman ρ [RPF]: mean 0.784 vs. ρ [RNA]: mean 0.736; p<2.2e-16) (Figure 2C; Figure S3C-D). This phenomenon was observed irrespective of whether the genes expressed only one dominant protein-coding transcript isoform (“single isoform genes” in the following) that was common to both organs, or whether they gave rise to different (including tissue-specific) mRNA variants (“multiple isoform genes”) (Figure 2D). The observed higher cross-organ concordance of RPFs could have simply had technical reasons, for example if the RPF-seq protocol gave more reproducible results than the RNA-seq protocol. We addressed this caveat by comparing measurement errors (MEs) for RNA and RPF data using a similar approach as in a recent publication [18]. We found that MEs scaled inversely with expression levels, as expected, and showed some variation due to organ (Figure S6A,B,F,G). Especially in liver and among low expressed transcripts, a tendency towards smaller MEs for RPF than for RNA was indeed visible (differences statistically non-significant). In most other cases, however, measurement errors were (in part significantly) higher for the transcripts’ RPF counts than for their RNA counts. Of note, the better cross-organ correlation of RPF vs. RNA levels seen in the full transcript set (Figure 2C) was also evident within various transcript subsets (Figure S6C-E, H-L), including such subsets for which RPF MEs were higher than RNA MEs (Figure S6E, L). It is thus unlikely that technical bias was the reason for the higher RPF correlation. Finally, an analysis that we performed on independent ribosome profiling datasets from rat liver and heart [19] allowed us to confirm the phenomenon of higher concordance of RPF vs. RNA abundance also between these organs(Figure S7A-C). Taken together, these findings are suggestive of a potentially broader biological phenomenon that consists in the partial compensation of differences in a gene’s mRNA expression through counteracting effects exerted through its TE, resulting in the convergence at the level of protein biosynthetic output (footprints, RPF) across tissues.

### Transcript features associated with cross-organ differences in translation efficiency

Do particular transcript features have predictive value for organ-specific differences in translation efficiency? To investigate this question we selected the genes with significantly different TEs between tissues (N=5013; Wilcoxon signed rank test; FDR<0.01) and implemented a 1.5-fold cutoff on TE ratio between the organs to retrieve the most pronounced cases (N=960) (Figure 2E; Supplementary Table S2). Of these, 533 represented “single isoform genes” with no (or negligible amounts of) expression of tissue-specific mRNA variants. For these genes, we examined whether a higher TE in kidney (N=193) or in liver (N=340) was associated with specific transcript characteristics. Of several features tested, we found that CDS and transcript lengths showed the most significant association with differential TE (Figure S8A-B). Of note, we had previously seen in liver that shorter coding sequences – i.e., transcripts encoding smaller proteins – are more efficiently translated [10]. Our present analyses suggest that such transcripts are also more prone to tissue-specific regulation at the translational level. Other sequence features showed some bias within the differential TE gene sets as well, although the effects were overall weaker and less consistent. Briefly, the 5′ UTRs of genes with higher TEs in liver were longer and predicted to fold more strongly. By contrast, transcripts with higher kidney TEs were associated with lower 5′ UTR GC content and slightly shorter 3′ UTRs. No association with differential TE was found for the Kozak sequence context score.

We also investigated two functional classes of sequence features, miRNA binding sites and upstream open reading frames (uORFs), for association with differential TE. Of note, the 960 “TE different” transcripts were not enriched for any predicted miRNA binding sites, making it unlikely that this class of post-transcriptional regulators is a major player in establishing tissue-specific TEs (data not shown). We had previously observed that in the liver the presence of a translated uORF in the 5′ UTR was strongly predictive of low TE at the main ORF [10]. An analogous relationship was evident in kidney as well (Figure S5C). To assess whether uORF translation was associated with TE differences across organs, we compared how the identified uORF-containing transcripts (i.e., single isoform genes showing translated uORFs in at least one organ; N=1377) distributed to the differential vs. non-differential TE gene sets. The group of genes with higher TE in liver was significantly enriched for transcripts with translated uORFs (p=6.08e-04; Fisher’s exact test) and there was slight depletion among genes with higher TE in kidney (not significant) (Figure S8C). Only few differential TE genes exhibited uORF translation that was exclusive to one organ, but there was a tendency for kidney-specific translation of uORFs to be associated with higher TE on the CDS in liver, and vice versa (Figure S8D). For the genes with uORFs translated in both tissues, we expected that cross-organ differences in the strength of uORF usage would negatively correlate with TE differences at the CDS. However, such a trend was only visible for liver differential TE genes (Figure S8E), and globally, uORF and CDS TEs even showed slightly positive correlation. In summary, these analyses suggested that uORF translation contributed to some extent (and especially for genes that were more efficiently translated in the liver) to cross-organ differences in TE; however, the overall impact appeared limited (see Discussion).

We next included the “multiple isoform genes” in the analyses and asked whether transcript isoform diversity between the two organs – i.e., the occurrence of tissue-specific mRNA variants generated by alternative transcriptional start sites, splicing, and 3′ processing – had any relationship to differential TE. Briefly, using our RNA-seq data we first compiled an inventory of all annotated, protein-coding transcript isoforms and their estimated relative expression levels per gene and tissue. We then used the Hellinger distance [20] as a measure of dissimilarity in isoform expression levels between kidney and liver. A value of 0 for this metric indicates that a gene has identical isoform distribution in both tissues (i.e., these are essentially the “single isoform genes” described above), while a value of 1 denotes a lack of overlap in expressed isoforms. Globally, the 960 genes with differential TE showed significantly higher Hellinger distances than the remainder of the expressed genes (p=3.74e-04; Kolmogorov-Smirnov-test) (Figure 2F). Molecularly, the term “transcript isoform” comprises variations affecting 5′ UTR, CDS, and 3′ UTR. By comparing the genes for which all expressed variants affected exclusively one single feature, or for which this particular feature was not affected at all, it became apparent that transcript diversity in the 5′ UTR was particularly strongly associated with differential TE (Figure 2G). By contrast, variation in the CDS showed significantly less association with cross-organ differences in translation efficiency (Figure S9A-B). Although the low number of available transcripts bearing exclusively 3′ UTR differences precluded a rigorous interpretation, 3′ UTR variation did not appear to be associated with differential TE either (Figure S9C). Altogether, we thus concluded that TE differences between tissues may, at least in part, have their origin in tissue-specific transcript variants, especially through alternative 5′ UTRs.

Finally, we were interested in whether cross-organ differences in translation efficiency affected specific pathways. For the 640 “TE different” genes that showed increased TE in liver (Figure 2D), gene ontology (GO) analyses revealed significant enrichment for categories related to transcription (Supplementary Table S2). Conceivably, tissue-specific translational control of transcriptional regulators may thus impact also on the organs’ transcriptomes. The 320 “TE different” genes that were translated better in kidney did not show any significant enrichment.

### Translational modulation of phase of oscillation in kidney

We next turned to the analysis of factor time across the datasets. We annotated rhythmic events in kidney with the same methodology as previously for liver, including a 1.5-fold cut-off on peak-to-trough amplitudes [10]. A list of the detected RNA and RPF rhythms and genome-wide gene expression plots are provided in Supplementary Table S3 and Supplementary Dataset S1, respectively. Our analyses yielded 1338 and 977 genes that cycled at the RNA abundance and footprint level, respectively, with an overlap of 542 genes (Figure 3A). As discussed later, this relatively modest overlap (542 genes corresponds to 41% and 55% of all “RNA rhythmic” and “footprints rhythmic” cases, respectively) likely underestimates the full extent of shared rhythmicity and only contains the most robustly oscillating gene expression events, which we further explored in the following.

**Fig. 3.**
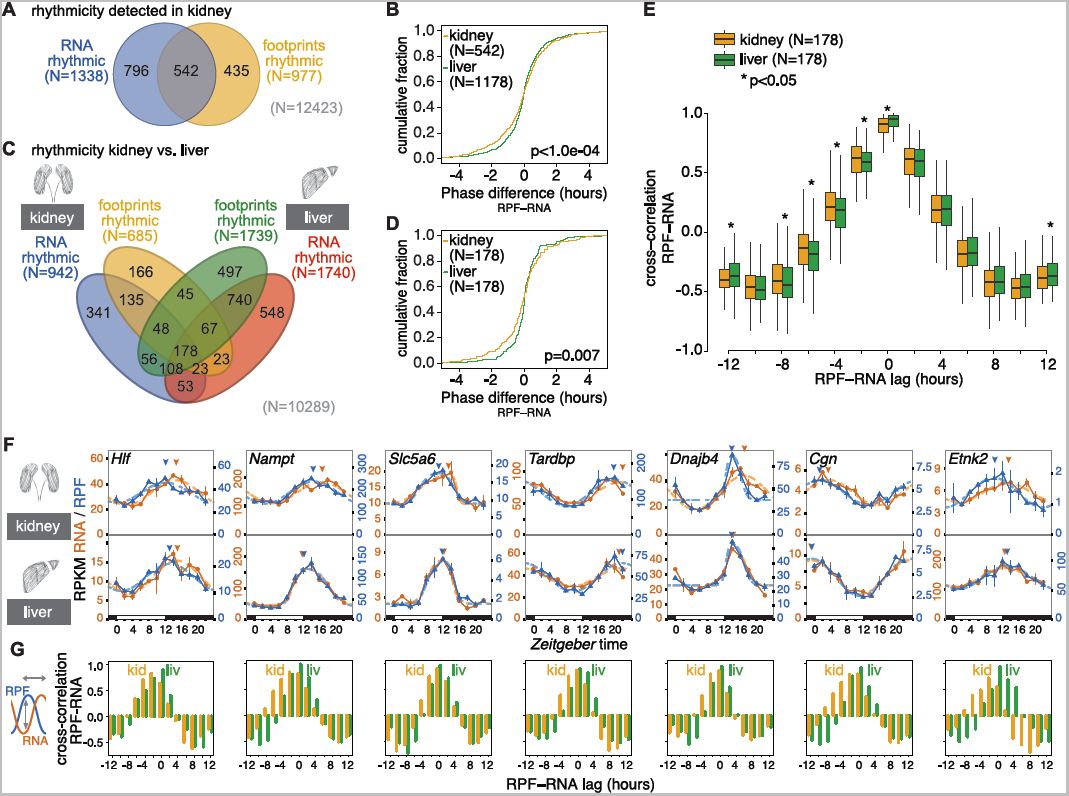
Rhythmicity analyses across organs reveals phase modulation by translation in kidney. **A** Venn diagram showing rhythmic genes in kidney. Of the 12423 expressed genes, 1338 showed 24-hour oscillations of >1.5-fold amplitude in mRNA abundance (RNA-seq, 10.7%) and 977 in footprint abundance (RPF-seq, 7.9%). 542 genes (4.3%) were identified as rhythmic at both levels. **B** Cumulative distribution of phase differences (RPF peak – RNA peak, in hours) for genesrhythmic at both RNA-seq and RPF-seq in liver (green, N=1178) and kidney (yellow, N=542). The two distributions were significantly different (p<1e-04, permutation test), and reflected that maximal footprint abundance frequently preceded mRNA abundance peaks in kidney (note that the two distributions differed mostly in their negative tail). **C** 4-way Venn diagram of rhythmicity sets for genes expressed in both tissues (N=10289). 364 and 238 genes were detected as rhythmic in both organs at the RNA-seq and RPF-seq levels, respectively, and 178 genes were detected as rhythmic throughout (i.e. RNA-seq and RPF-seq, in kidney and liver). **D** Cumulative phase difference distribution in liver (green) and kidney (yellow) for the 178 common rhythmic genes. As in (B), the distributions were significantly different (p=0.007, permutation test), and corroborated that even when comparing the same set of genes, footprint peaks frequently preceded mRNA abundance maxima in kidney. **E** Cross-correlation in kidney (yellow) and liver (green) of time-resolved RPF-seq profiles relative to the RNA-seq profiles of the N=178 common rhythmic genes. The analysis showed that profile correlations for negative lags (i.e. RPF peaking before RNA) were significantly higher in kidney than liver (asterisks indicate p<0.05, Wilcoxon signed rank test). Boxplots represent the interquartile range and whiskers extend to the minimum and maximum expression within 1.5 times the interquartile range. **F** Examples for genes with maxima in RPF (blue) preceding those in RNA (orange) by several hours in kidney (top) but not, or less so, in liver (bottom). Arrowheads indicate the peaks in footprint and mRNA abundance as estimated from the rhythmic fits. **G** Cross-correlation analyses of RPF-seq relative to RNA-seq profiles (kidney in yellow, liver in green) for the genes in (F). Maximal correlations of the profiles in kidney were found to be shifted to the left (more negative RPF-to-RNA lags) as compared to liver. For liver, there was no case with a maximal correlation value in the negative.

Interestingly, the analysis of rhythmicity parameters across the 542 genes revealed that the timing of their RPF peaks relative to their RNA peaks had a significantly different and broader distribution than the corresponding set from liver (p<1.0e-04; permutation test) (Figure 3B). This observation suggested that the phase of protein biosynthesis rhythms was subject to marked translational modulation in kidney. In liver, by contrast, RPF peaks were more tightly gated by RNA abundance peaks. Surprisingly, maximal translation tended to *precede* maximal RNA abundance in kidney (Figure S10A), as globally the mean RPF peak phase was advanced (-0.123 hours) and also RPF rhythms were enriched for phase advances (282) vs. delays (260), albeit neither reaching statistical significance (p=0.16, Wilcoxon rank sum test).

The above analyses used different rhythmic gene sets for kidney than for liver, potentially compromising comparability. The observed differences in the RPF-RNA phase relationships could thus have simply arisen from transcript-specific rather than from tissue-specific differences in the timing of translation. We thus analysed the group of 178 genes whose RNA and RPF profiles were rhythmic in both organs (Figure 3C; Supplementary Table S4; Supplementary Dataset S2). Again, the distribution of RPF-RNA offsets was significantly broader in kidney than in liver (Figure 3D; p=0.007, permutation test) with an RPF peak phase advance in kidney (mean -0.143 h) and a phase delay in liver (mean 0.036 h) (Figure S10B-C). We next calculated the gene-wise RPF-RNA peak phase difference in kidney relative to that in liver. More genes showed their RPF maxima earlier (96) than later (82) in kidney vs. liver, with a mean advance of -0.178 h (Figure S10D), but again without passing statistical significance (p=0.152, Wilcoxon rank sum test).

Conceivably, we lost statistical power and introduced error in the above analyses by restricting the phase comparisons merely to the peaks of the rhythmic curve fits. We thus sought a method that would take into account phase differences between RPF and RNA profiles over all data points. To this end, we used cross-correlation to quantify the similarity between the RPF and RNA time series as a function of sliding one series on the time axis relative to the other. When the time series were not shifted against each other at all (RPF-RNA lag = 0 h), the RPF-RNA cross-correlation values were overall highest, as expected, and they were significantly higher in liver, in line with stronger gating of RPF rhythms relative to RNA oscillations in this organ (Figure 3E). Importantly, when cross-correlation of RNA was calculated with earlier RPF time points (negative RPF-RNA lags; see in particular lags of -2 h to -8 h in Figure 3E), kidneys scored significantly higher than livers. Sliding the series in the other direction, however, rather led to overall better correlations in the liver (see lags of +4 to +8 in Figure 3E; liver-kidney difference non-significant). Taken together, these analyses underscored that there was asymmetry in the data with RPF rhythms preceding RNA rhythms specifically in the kidney.

We confirmed kidney-specific translational phase advances by visual inspection of individual gene expression profiles. Figure 3F shows the profiles for the genes *Hlf, Nampt, Slc5a6*, *Tardbp*, *Dnajb4*, *Cgn* and *Etnk2*, which all show an RPF phase advance of up to several hours relative to RNA. Cross-correlation analysis for the individual genes confirmed kidney-specific, phase-advanced translation as well (Figure 3G).

At first sight, translation that is phase-advanced to mRNA abundance is counterintuitive. Conceivably, it may occur when translation efficiency is not constant, but decreases over the lifetime of an mRNA. TEs may be higher on freshly synthesised messages that have long poly(A) tails, and decrease as a result of gradual deadenylation even before transcript stability and abundance are affected as well [21]. In keeping with the hypothesis of cross-organ differences in poly(A) kinetics, we have observed that most subunits of the major cytoplasmic deadenylase complex, CCR4-NOT, are significantly more highly expressed in kidney than in liver (Figure S11A-C). Higher deadenylase activity in kidney could provide an attractive molecular explanation for the observed tissue-specific differences in RPF-RNA phasing, and for RPF rhythms that are phase-advanced to RNA oscillations.

### High tissue divergence in translationally driven rhythms

Rhythmicity detection algorithms are sensitive to false-negatives i.e., to classify gene expression profiles as “non-rhythmic” (for example because they fail imposed thresholds on amplitude or FDR) although the underlying temporal patterns may still be more similar to, and more likely to be, rhythmic than invariable. Of note, the lack of canonical methods to reliably determine true absence of rhythms is a common problem in the field (see recent review by [6] for discussion). Venn diagrams that simply overlap rhythmic gene sets hence need to be interpreted with caution. For these reasons, the extent of “RNA only” and of “footprints only” oscillations in Figure 3A is likely not reported reliably and subject to overestimation. The heatmaps of the corresponding RNA and RPF profiles support this notion as well (Figure S12B, D).

In order to identify the true-positive “translation only” cycling transcripts with higher reliability, we implemented the same methodology as in our previous study [10]. Briefly, we used the analytical framework *Babel* [22] to preselect all transcripts whose translation efficiency changed significantly over the day (and/or whose TEs deviated significantly from the global transcript population). Rhythmicity analyses were then performed on this gene subset and yielded 92 cases with the sought-after temporal profiles of rhythmic translation on non-rhythmic mRNAs (Figure 4A). Comparison with the 142 genes of the analogous set from liver revealed near-perfect tissue specificity of translationally driven oscillations. Only two genes, *Abcd4* and *Lypla2*, were shared between the organs; they were both among the least compelling cases of “translation only rhythms” that our method had identified, as judged by visual inspection (Figure 4B).

**Fig. 4.**
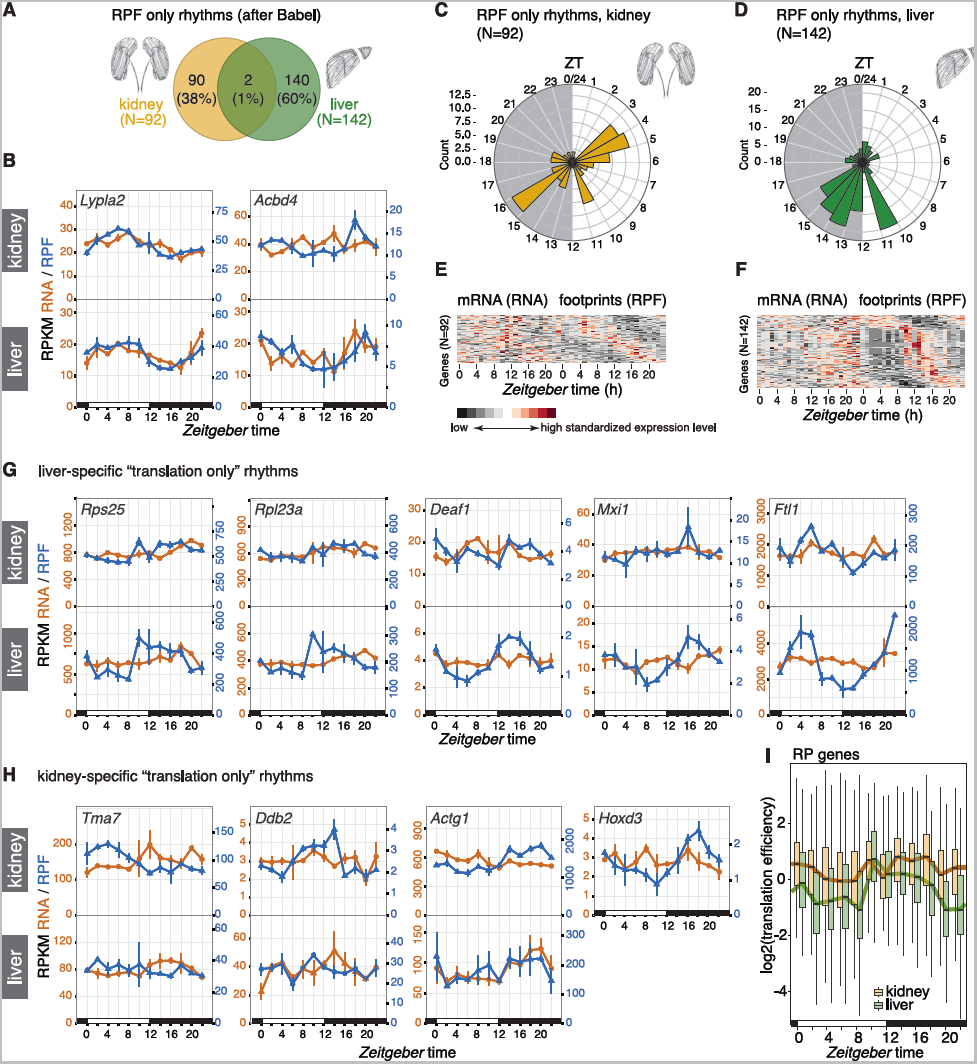
High tissue divergence in translationally driven rhythms. **A** Venn diagram of rhythmic RPF-seq sets in kidney (yellow, N=92) and liver (green, N=142) afterthe Babel analysis indicated strong tissue-specificity of translational control. **B** Daily profiles of RPF-seq RPKM (blue) and RNA-seq RPKM (orange) for the two genes detected as translationally regulated in both tissues in A. **C** and **D** Circular phase histogram for the 92 (C, kidney) and 142 (D, liver) genes showing footprint rhythmicity in the organs. Note that the translational upregulation of transcripts observed at the day-to-night transition in liver was absent in kidney. **E** and **F** Heatmaps of RNA (left panels) and RPF (right panels) rhythms for the 92 and 142 genes translationally regulated in kidney (E) and in liver (F), respectively. Genes are sorted by footprint phase and expression levels are standardized by row (gene). These sets of genes showed rhythmicity in footprint abundance but no oscillation in mRNA. **G** and **H** Daily profiles of RPF-seq RPKM (blue) and RNA-seq RPKM (orange) for representative examples of translationally generated rhythms specific for liver (G) and kidney (H). For each gene, the upper panel shows the kidney data and the lower panel the liver data. *Hoxd3* was not expressed in liver. **I** Translation efficiency (TE) around-the-clock for ribosomal protein (RP) genes expressed in liver (green, N=86) and in kidney (yellow, N=89). For each timepoint (ZT) boxplots represent the interquartile range and whiskers extend to the minimum and maximum TE within 1.5 times the interquantile range. Lines connect the median of each boxplot to ease visualization. Note the global TE upregulation at ZT10 in liver, whereas TEs in kidney remain high over the day.

Interestingly, not only the identity of rhythmically translated genes, but also the time-of-day at which the majority of rhythmic translation events occurred, was highly tissue-specific. The phase histograms thus showed striking differences in the peak time distributions between the organs (Figure 4C-D; difference in distributions: p=1.66e-04; W = 17.403, df = 2; Watson-Wheeler test for homogeneity of angles). Of note, the enrichment for translational maxima at the light-dark transition (*Zeitgeber* time, ZT10-16; ZT00 corresponds to lights-on and ZT12 to lights-off) that dominated the distribution in liver (Figure 4D, F) was virtually absent from kidney (Figure 4C, E). Instead, kidney showed enrichment for transcripts with maximal translation occurring around ZT4 and ZT16. Visual inspection of individual examples confirmed the organ specificity of RPF rhythms. The cases of robust translational oscillations that we [10] and others [11, 12] had previously identified in liver were thus absent or severely blunted in kidney. This included mRNAs encoding ribosomal proteins (RPs), which make up the bulk of genes showing a translational surge at the light-dark transition (e.g. *Rps25*, *Rpl23a*), as well as transcripts encoding the transcription factors *Deaf1* (deformed epidermal autoregulatory factor 1) and *Mxi1* (MAX interactor 1), and mRNAs containing iron-responsive elements in their 5′ UTRs (e.g. Ferritin light chain 1, *Ftl1*) (Figure 4G), all of which we had previously reported as translationally rhythmic in liver [10]. Rhythmic translation exclusive to kidney was not significantly enriched for particular pathways (data not shown), and the temporal profiles were overall of lower amplitude than those seen for liver; *Tma7* (Translational machinery associated 7 homolog), *Ddb2* (Damage-specific DNA binding protein 2), *Actg1* (Actin, gamma, cytoplasmic 1) and *Hoxd3* (Homeobox D3; not expressed in liver) were among the most distinct examples (Figure 4H).

In summary, we concluded that temporal changes in TE were strikingly tissue-specific and overall relatively rare in kidney. Specifically for transcripts encoding RPs and other components of the translation machinery, which are the most prominent group of TE rhythmic genes in liver, it has been suggested that feeding-dependent mTOR-signaling underlies the translational upsurge at the light-dark transition via a mechanism involving the 5′-terminal oligopyrimidine (5′-TOP) motifs that these transcripts carry [11, 12]. Interestingly, the TE comparison between both tissues revealed that kidney RP translation occurred at a relatively high level throughout the day (Figure 4I). The lack in rhythmicity for RP genes in this organ may thus result from an absence of translational repression during the light phase rather than a lack in activation in the dark phase. It may indicate that the kidney is less sensitive to systemic cues engendered by feeding and fasting (see Discussion).

### Different degrees of tissue specificity in core clock gene expression at the level of RNA abundance and protein biosynthesis

Clocks exhibit functional differences across cell types and organs, for example at the level of rhythmicity parameters (e.g. free-running period and phase [5]), of clock output gene repertoires [3], of oscillator strength and robustness [23, 24], or with regard to clock gene loss-of-function phenotypes [25]. Conceivably, the precise timing and level at which the various clock proteins are produced may modulate properties of the clock circuitry and underlie some of the abovementioned functional variations. In order to investigate these possibilities we compared the expression of core clock components in both organs.

We first investigated transcript and footprint RPKMs as averages over timepoints to assess the cumulative daily production of clock RNAs and proteins. Most core clock genes showed a considerable degree of organ-specificity in their expression levels that was readily appreciable in the footprint vs. transcript abundance representation with both organs overlaid in a single graph (Figure 5A). Two tissue differences caught our particular attention. First, the balance between the transcriptional activators *Rora/Rorc* and repressors *Nr1d1/Nr1d2* differed markedly between organs and was skewed towards repression in kidney (i.e., higher *Nr1d1/2* and lower *Rora/c* RPKMs in kidney, Figure 5A-B). These transcriptional regulators bind to shared sequence elements on DNA and form the “interconnecting limb” within the rhythm-generating clock circuitry. In addition, they also control an output branch of the oscillator [1, 2]. It is hence conceivable that adjusting the relative levels of NR1D1/2 vs. RORs tailors clock-controlled gene expression in a tissue-specific fashion. Our observation of an active state of this output branch in liver and a more repressed state in kidney, is fully consistent with the current knowledge of its target genes and knockout phenotypes, which point to a prominent role in the regulation of hepatic pathways such as lipid, cholesterol and bile acid metabolism [26].

**Figure 5.**
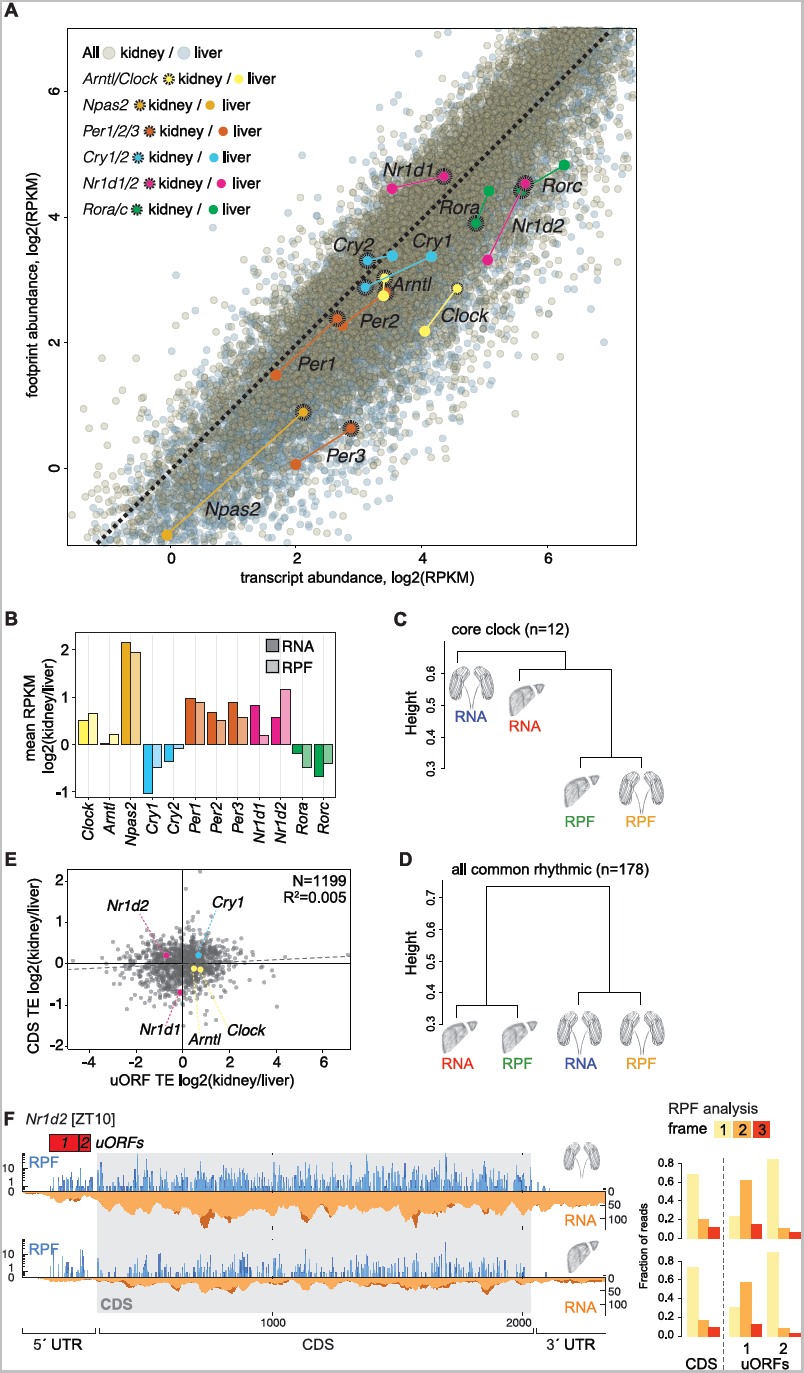
Tissue specificity in core clock gene expression at the level of RNA abundance and translation. **A** Scatterplot of transcript abundance (RNA-seq) vs. footprint abundance (RPF-seq) for liver (grey) and kidney (sepia) (N=10289), where core clock components are highlighted (kidney dots with dashed circles). Coloured dashed lines join the relative locations of each core clock gene between organs. **B** Bar graph of the average RPKM ratio between kidney and liver for the main circadian core clock genes, at the level of mRNA abundance (dark shades) and ribosome footprints (light shades) suggested that translational compensation led to higher similarity at the level of protein biosynthesis (RPF) for several core clock genes. **C** Hierarchical clustering of the organs’ RNA and RPF profiles based on the similarities of the core clock genes expression patterns (N=12, genes shown in B). The height of the branches represents weighted average distances over the considered genes (see Methods). Note that RPF rhythms in two organs were more similar than RNA and RPF rhythms within an organ. **D** Hierarchical clustering as in C based on the genes detected as rhythmic throughout (N=178, see Figure 3C). When compared to the clustering based on core clock gene expression patterns in C, this rhythmic gene set showed an organ-based clustering. **E** Scatterplot of kidney/liver ratios of uORF vs. CDS translation efficiencies for genes containing AUG-initiated translated uORFs in both organs (N=1199). uORF-containing core clock genes are highlighted. As also shown in Fig S8E, differential uORF usage could not globally explain differences in CDS TE across organs (note the lack of negative correlation between the two variables, R^2^ = 0.005, p=0.008). As an exception, the lower uORF TE of *Nr1d2* might have a role in setting relatively higher CDS TE in kidney. **F** RPF (blue) and RNA (orange) reads mapping along the *Nr1d2* transcript in kidney (top) and liver (bottom) for the timepoint of maximal CDS translation (ZT10). 5′ UTR and CDS are shown in full, but for better visualization only a portion of the 3′ UTR (the same length as the 5′ UTR) is shown. Red boxes indicate the predicted AUG-initiated translated uORFs. Right panels show that, similar to the CDS, the uORFs showed clear frame preference, indicative of active translation.

A second tissue difference concerned the main constituents of the negative limb, the Period (*Per*) and Cryptochrome (*Cry*) genes. Heterotypic PER:CRY protein complexes inhibit ARNTL:CLOCK-driven transcriptional activity and thus lie at the core of the oscillator’s principal negative feedback loop. We observed a shift to more PER (and slightly less CRY) biosynthesis in kidney (Figure 5B). PERs (and in particular PER2) are considered stoichiometrically rate-limiting components of the PER:CRY complex, and increased PER2 dosage engenders long periods [27, 28]. Interestingly, tissue explant experiments have shown that kidney clocks free-run with almost 1.5 hour longer periods than liver clocks [5], as would be predicted from the increased PER biosynthesis that our analyses revealed. As a more general concept, we deem it conceivable that the modulation of biosynthesis levels for individual clock proteins may be a more general mechanism to engender distinct differences in clock parameters across cell types.

We noted that for the majority of clock genes (*Npas2, Cry1, Cry2, Per1, Per2, Per3, Nr1d1, Rorc)* the tissue differences were less pronounced at the RPF than at the RNA level (Figure 5B), indicating that translation efficiencies partially counteracted RNA expression differences. Only in four cases (*Clock, Arntl, Nr1d2, Rora*) TEs exacerbated transcript abundance differences and led to higher tissue differences at the RPF level. Interestingly, this observation could also be made in the time-resolved data. As a measure of similarity between rhythmic profiles, we used the Euclidean distances calculated between the four rhythmic traces of each individual gene (i.e., RNA and RPF in kidney and liver; Figure S13). Hierarchical clustering of the similarities for the ensemble of the 12 main core clock genes showed that RPF rhythms from the two organs grouped together (Figure 5C). Clock protein biosynthesis rhythms between organs were thus more similar than RNA and RPF rhythms within organs. By contrast, the 178 common rhythmic genes identified in Figure 3C – serving as a control set for this analysis – revealed within-organ clustering (Figure 5D). These findings underscored that translational compensation was occurring within the core clock, where it led to more similar rhythms in clock protein biosynthesis than would have been predicted from the rhythmic RNA abundance profiles. This phenomenon was, however, not a general feature of all rhythmic gene expression.

The transcriptome-wide analyses described further above had shown only weak signs of association between cross-organ differences in TE and in uORF usage (Figure S8C-E). However, we knew from our previous work in liver that at least five core clock transcripts (*Nr1d1, Nr1d2, Cry1, Clock, Arntl)* contained translated, potentially regulatory, AUG-initiated uORFs [10]. We therefore examined whether for any of these concrete cases there was evidence for a connection between uORF translation and cross-organ TE differences. The read distribution along the transcripts (Figure S14A) and the marked frame preference of RPF reads (Figure S14B) confirmed that the footprints mapping to our annotated uORFs likely reflected active translation. However, only in one case, *Nr1d2*, there was a distinct anticorrelation between uORF usage and TE differences on the CDS (Figure 5E). *Nr1d2* contains two translated uORFs in the 5′ UTR (Figure 5F), whose decreased usage in kidney was accompanied with higher TE on the CDS in this organ (Figure 5E). For *Nr1d2*, differential uORF usage could thus represent a plausible mechanism that contributes to regulating organ-specific gene expression output at the translational level, keeping NR1D2 biosynthesis low in liver and high in kidney.

## Discussion

Given that the functionally relevant output of most gene expression is the protein, quantitative and genome-wide analyses of protein biosynthesis are of high interest to complement the wealth of transcriptomics data that is already available. Of note, only the fairly recent development of the ribosome profiling technique [29] has made it possible to analyse translational events in a quantitative, high-throughput fashion. Our study of two paradigms of differential gene expression, i.e. its tissue- and its time of day-dependence, is among the first of its kind and, together with the associated datasets and resources, will likely be of wide interest and utility to researchers working in the chronobiology and gene expression fields.

We have addressed several, rather fundamental questions that go beyond the chronobiological focus of the study: How does the dynamic range of translation efficiency compare to that of transcript abundances across two distinct organs of an animal? Is translation efficiency a default transcript property and comparable across two tissues, or do TEs become reinterpreted depending on cell type or organ? Does cross-tissue variability of TEs come with any direction i.e., is there a global tendency to either reinforce or to counteract transcriptomal differences?

To our knowledge, only one previous study has reported on ribosome profiling datasets from two complementary mammalian tissues, rat liver and heart [19]. This study also included animals with different genetic backgrounds as covariates in the experimental design, and its main focus was on strain differences in translation rather than on tissue differences. Our analyses based on more than 10000 genes commonly expressed in liver and kidney show that cross-organ TE differences are widespread, but of limited magnitude. Across genes in a tissue and for individual genes between tissues, the dynamic range of translation efficiencies is thus about 30-50-fold narrower than that of transcript abundances. These findings are coherent with the view that major differences in gene expression are set up at the level of transcription (possibly with some influence coming from RNA stability as well), whereas differences in translation rate have more of a modulatory role. It is intriguing that this modulation is overall characterised by directionality, with TE differences between tissues globally counteracting some of the mRNA abundance differences. Such translational compensation has previously been observed for divergent transcript expression levels across yeast species [30] and across different rat strains [19]; our study now extends this observation to gene expression across organs. Moreover, the idea of translational compensation is conceptually similar to findings that proteomes are evolutionarily more highly conserved than transcriptomes [31, 32]. As an underlying common principle, these cases may indicate that selective pressure on precise gene expression levels likely acts on protein abundances, whereas a certain degree of variability (even noise) in RNA levels may be tolerated without further consequences. It will be exciting to study the underpinnings of translational compensation further, across tissues and across species.

Maybe not unexpectedly, there was no dominant, distinct sequence feature that could serve as a predictor for cross-organ TE differences. Rather, we found several associations with a number of transcript characteristics. Conceivably, these contribute collectively to modulating TEs in concert with the specific cellular and tissue environment and possible cell type differences in the translation machinery including its regulators and trans-acting factors. While our ribosome profiling studies have allowed us to record the outcome of such regulation at high resolution, understanding its causes represents an exciting challenge for the future. For now, we can only infer that an overarching theme of the identified associations is a connection to 5′ UTRs, which is in also in line with the notion that initiation is rate-limiting for most translation events. We thus observed associations of cross-organ TE differences with 5′ UTR length, with uORF usage, with GC content and folding potential, as well as with transcript isoform diversity that affected the 5′ UTR. We would like to point out that comprehensive uORF annotations remain a bioinformatics challenge that is far from resolved. We have therefore restricted our analyses to AUG-initiated ORFs, inevitably leading to a bias towards false-negatives in uORF annotation. As we will learn how to annotate uORFs more comprehensively and more precisely in the future, it may be worth revisiting the relationship between differential TE and uORF translation in our datasets in order to evaluate whether a clearer role for these regulatory sequence elements will emerge.

Our study has led to novel insights into rhythmic gene expression. The extent to which rhythmicity is generated by the temporal regulation of translation has been the subject of speculation ever since the first report of rhythmic proteins encoded by non-rhythmic mRNAs [7]. Our kidney datasets complement recent time-resolved ribosome profiling data from liver [10, 11] and from a cell line [33]. As compared to liver, the number of transcripts subject to translational rhythms in kidney is slightly lower, but overall in a similar order of magnitude with around 1% of the transcriptome affected. It came as a surprise that translational rhythms were essentially tissue-specific in terms of the affected genes and the phase distributions. A possible explanation could be that these rhythms are driven by rhythmic systemic cues to which tissues do not respond equally. The effects of feeding and mTOR signalling, for example, may be more pronounced in liver due to the dedicated role that this organ plays in energy homeostasis and fasting responses, thus explaining the differences in translational oscillations for RP genes. Beyond the role that translation has in generating rhythms, our analyses have pointed to an additional, rhythmicity-modulating role that appears to affect gene expression quite broadly, i.e. the timing of the phase of protein biosynthesis oscillations relative to that of mRNA abundance rhythms. Consistent with work by the Green lab that showed interactions between polyadenylation status of mRNAs and rhythmic protein expression in the liver [21], it is tempting to speculate that related mechanisms are operative across organs, with tissue-specific deadenylation kinetics tuning the timing of rhythmic protein biosynthesis. Finally, our study is a first attempt to resolve tissue differences in core clock gene expression as a factor contributing to functional differences of the oscillator. It is interesting that the core clock mechanism has a long-standing history of being referred to as a “transcription-translation feedback loop” [34], although the actual feedback occurs at the transcriptional level, and possible mechanistic functions of translational regulation have not been much investigated. Our cross-organ comparison of core clock protein biosynthesis suggests that translational control – including through the activity of uORFs [10, 33] – is of regulatory interest and represents a way by which the identical set of core clock genes could form circuitries with different stoichiometry of its main components. As a result, both clock parameters and output gene repertoires may be organ-specifically tuned.

## Conclusions

How translational differences contribute to overall gene expression diversity is still poorly understood. Our study uncovered translational changes that occur across two paradigms of regulated gene expression, i.e. around-the-clock and between tissues. Daily gene expression rhythms generated at the translational level were strongly organ-specific with regard to the identities and phase distributions of affected genes. Moreover, our data indicate that translation efficiency differences between organs can adjust the timing of protein production from rhythmic mRNAs, and the levels of core clock protein production, in agreement with the tissue-specificities observed in clock output gene sets and clock parameters. Together, these results are consistent with an important role of post-transcriptional mechanisms in mammalian circadian gene expression regulation. Beyond the temporal dimension, we have explored constitutive protein biosynthesis across organs. Our quantitative analyses underscore that gene expression divergence is largely programmed at the transcript abundance level. Interestingly, the widespread differences in translational efficiency that we detected between organs even serve to achieve higher concordance in protein production between tissues. Conceivably, such translational compensation reflects a selective pressure to maintain precise protein levels rather than mRNA levels. The high-resolution genome-wide translatome datasets generated in this study will allow further explorations into the mechanisms of post-transcriptional control and differential gene expression *in vivo*.

## Methods

### Animals

12-week old male mice (C57BL/6J; Janvier Labs) were entrained for two weeks to light:dark 12:12 cycles with *ad libitum* access to food and water and were anesthetized (isoflurane) and sacrificed every two hours (ZT0 - ZT22, with ZT0 corresponding to “lights-on”) for two daily cycles. Livers and kidneys were removed and processed either directly or flash-frozen in liquid N_2_. All experimental procedures were approved by the Veterinary Office of the Canton Vaud (authorisation VD2376).

### Ribosome profiling

Generation of liver RPF-seq and RNA-seq libraries using the ARTseq ribosome profiling kit (Epicentre) was described recently [10, 16]. Kidney libraries were prepared in the same manner, with a single modification to the order of steps in RPF library preparation. After RNase treatment and recovery of ribosome-protected fragments, 5μg of material was first rRNA-depleted (Ribo-Zero magnetic kit, Epicentre) and then purified by 15% PAGE. In the formerly prepared liver libraries, Ribo-Zero treatment and PAGE purification had been inverted because at the time we had found that changing the order had a beneficial effect on obtaining highly concentrated libraries. For the kidney samples, however, we noted that this modified order led to higher contamination with reverse-strand rRNA probes bleeding from the Ribo-Zero kit and we thus reverted to ARTseq’s original order. All other steps and materials were identical between liver and kidney samples and followed the ARTseq ribosome profiling kit instructions. RPF and RNA libraries were sequenced on an Illumina HiSeq 2500.

### Sequencing data processing, alignment and quantification

Processing, quality assessment, alignment and quantification of sequencing data were performed as described previously [10, 16]. Briefly, after adapter trimming using Cutadapt [35], the length distribution of trimmed reads was used to assess the quality of nuclease digestion and size-selection, which is particularly important for RPF libraries (Figure S1B). Trimmed reads were filtered by size (26-35 nt for RPF; 21-60 nt for RNA) using an in-house Python script, and sequentially mapped to mouse rRNA, human rRNA, mitochondrial tRNA, mouse tRNA, mouse cDNA (Ensemble mouse database release 75) using Bowtie v2.2.1 [36] and mouse genome (GRCm38.p2) using Tophat v2.0.11 [37]. Trimmed and filtered sequences were also directly mapped against the mouse genome (Tophat v2.0.11) in order to estimate expressed transcript models in each organ (using Cufflinks v2.2.1 [38]). Transcriptome-mapping reads in the sequential alignment were counted towards their location into the 5′ UTR, CDS or 3′ UTR of the transcript, based on feature annotation (Ensemble mouse release 75). Mappable and countable feature lengths were not calculated for this study (see “faux reads analysis” in the “Quantification of mRNA and ribosome footprint abundance” section of Supplemental Experimental Procedures of previous study [10]) as its contribution was negligible for further analyses. Therefore RPKM calculations in this study were not corrected with such factor. Read counts in RNA-seq and RPF-seq datasets were normalised with upper quantile method of edgeR [39] and RPKM values were calculated as the number of reads per 1000 bases per geometric mean of normalised read counts per million. Relative translation efficiencies (TE) were calculated as the ratio of RPF-RPKM to RNA-RPKM per gene per sample. Reading frame and nucleotide periodicity analyses were performed as in [10]. Principal Component Analysis (PCA) relied on a combined matrix of CDS counts for RPF and RNA from both liver and kidney and following the same approach as before [10]. Ribo-seq Unit Step Transformation (RUST) analysis was used to assess whether the sequencing libraries were globally of similar quality in terms of their local footprint densities [17]. RUST is a simple normalisation method that reduces the heterogeneous noise in the data and allows identification of mRNA sequence features that affect footprint densities globally. We used the version 1.2 of the published rust_codon.py standalone python script with minor modifications to reflect the experimental settings as closely as possible (i.e., A-site offsetting). RUST codon profile and corresponding Kullback–Leibler (K–L) divergence for each library (RPF and RNA) was generated against a database of 8012 single protein isoform transcript sequences using all mapped reads with a length of 28-32 nt. The K-L divergences from all samples for each combination of tissue (kidney or liver) and read type (RPF or RNA) were used to generate K-L profiles at the 0, 10, 25, 50, 75, 90, and 100th quantiles.

### Correlation analyses and assessment of translational compensation across organs

Correlation of RNA-seq and RPF-seq across organs: Kidney vs. liver correlations at the levels of RNA- and RPF-seq (i.e., Figure 2B; Figure S6C-E, H-L) were calculated in a pairwise fashion for each of the 24 samples (12 timepoints, 2 replicates/timepoint), as livers and kidneys of each replicate originated from the same animals. Significance of the difference in the Spearman coefficients between both distributions was assessed by paired t-test on Fisher z-transformed coefficients. Heart vs. liver correlation at the levels of RNA and RPF-seq (Figure S7A) was calculated from the study [19], using the BN-Lx reference rat strain data. Since the five heart and liver replicates in this study did not come from the same animals, we calculated all possible pairwise correlation coefficients between heart and liver (i.e., 25), and compared all possible combinations of 5 coefficients between RNA- and RPF-seq (paired t-test on Fisher z-transformed coefficients).

Measurement error: Measurement errors (Figure S6A,B,F,G) were calculated similarly to [18] using the meas.est() function from smatr R package [40]. Genes were first binned according to average expression level (calculated as the fourth root of the product of liver RNA-seq, liver RPF-seq, kidney RNA-seq and kidney RPF-seq) into 10 groups, each containing 10% of all genes. Within each bin, the measurement error was calculated separately for RNA- and RPF-seq and for liver and kidney, using the two replicates (log of normalised CDS counts) to estimate the error, and the 12 timepoint samples to estimate its variability. For the analyses using a filtered gene set (Figure S6F-G), genes that showed a mean expression ratio (either between organs or between RNA- and RPF-seq) greater than 2 for all timepoints were excluded (9236 genes used in analysis).

### Analyses of differential translation efficiency

To test for differential translation efficiency (TE) between liver and kidney we used the Wilcoxon-signed rank paired test, using all 24 samples (12 timepoints; 2 replicates/timepoint) as replicates; resulting p-values were FDR-corrected. A gene was defined as having differential TE when FDR < 0.01 and the inter-organ difference in TE was at least 1.5-fold (Figure 2E).

Analysis of transcript usage diversity across organs: For each gene g, P(g) = (p 1, …, p n) is the vector of the relative expression proportions of its n protein-coding transcripts, as estimated from our RNA-seq analysis (see Sequencing data processing, alignment and quantification). To quantify the dissimilarity in relative transcript isoform expression between liver L and kidney K, the Hellinger distance H is defined as:

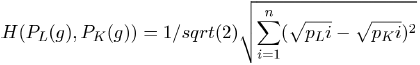

In order to detect the transcript features that were associated with tissue specificity in TE, we selected genes whose transcript diversity between both organs originated from, or was excluded from, 5′ UTR, CDS, or 3′ UTR, based on feature annotation information for the detected protein-coding transcripts (Figure 2G and Figure S9).

Study of transcript characteristics: For single-isoform genes, we investigated whether a particular transcript characteristic (length, GC content, Kozak context, structure) could be predictive of differential TE. Length and GC content were determined directly on the whole transcript and/or on the region of interest (5′ UTR, CDS, 3′ UTR). Kozak context was scored according to the consensus sequence GccA/Gcc**AUG**G, where upper-case letters denote highly conserved bases (scored +3), lower-case letters indicate the most common nucleotides (scored +1), and bold is the start codon (not scored), giving a maximum score of 13. Minimum free energy secondary structures on the 5′ UTR were predicted with RNAfold from ViennaRNA package with default parameters [41].

### Detection and translation efficiency calculation for uORFs

To assess the impact of differential uORF usage on TE differences across organs, uORFs were identified as in our previous study [10]. Briefly, genes expressing a single protein-coding isoform in both organs were used for this analysis (N=5815). We selected uORFs with an AUG start codon and a length of at least 18 nt to the first in-frame stop codon and considered them as translated if the reads showed significant frame bias towards the reading frame of the uORF start codon and if coverage was >10%. uORF translation efficiency was calculated from the ratio of RPF-seq to RNA-seq reads whose predicted A-sites mapped to the annotated uORF regions. If several uORFs partially of completely overlapped on a given 5′ UTR, a composite uORF was considered for read counting. uORFs overlapping with the CDS in the same frame were not considered. When they overlapped in different frames, only reads mapping to the 5′ UTR-specific uORF sequence (but not the overlapping sequence) was considered for quantifications.

### Rhythmicity analyses

Rhythmicity detection and rhythmic parameter estimations in each dataset (RNA-seq and RPF-seq, liver and kidney) were done based on Akaike information criterion (AIC) model selection as in our previous study [10]. The Babel computational framework [22] was used to detect rhythmically translated genes from constantly expressed mRNAs within each organ. For cross-correlation of time series to compare the daily profiles of rhythmic genes beyond their peak differences, we used the ccf function in R. As we computed the correlations of the RPF-seq with respect to the RNA-seq profiles, negative lag values correspond to RPF leading RNA.

### Hierarchical clustering of rhythmic genes

To evaluate the similarity of the expression profiles for rhythmic genes, a dissimilarity matrix was computed for each gene of interest, based on the Euclidean distance between the RNA-seq and RPF-seq expression profiles within and across organs. A hierarchical clustering tree was constructed on the weighted average of the dissimilarity matrices of genes under consideration (core clock genes in Figure 5C or all rhythmic genes in Figure 5D), using the “average” clustering method. The R functions{packages} dist{stats}, fuse{analogue} and hclust{stats} were used for computing the individual dissimilarity matrices, the weighted mean dissimilarity matrix, and the hierarchical clustering, respectively.

## Acknowledgments

We thank Lausanne Genomic Technologies Facility and Vital-IT staff for high-throughput sequencing and computational support. DG acknowledges support by: Swiss National Science Foundation (grants 128399, 157528); National Centre of Competence in Research (NCCR) RNA & Disease; Fondation Pierre Mercier; Fondation Leenaards; Olga Mayenfisch Stiftung; SystemsX.ch StoNets consortium; University of Lausanne. PJ was supported by Human Frontiers Science Program long-term fellowship LT000158/2013-L.

## Declarations

### Electronic supplementary material

CasteloSzekely_Suppl.pdf: This file contains the Supplementary Figures S1-S14

CasteloSzekely_Suppl_Table_S1.xls: This file contains information on the deep-sequencing data from kidney (raw read counts, mapping summary etc.)

CasteloSzekely_Suppl_Table_S2.xls: This file contains details of the GO-term analysis on the differential TE gene set of Figure 2.

CasteloSzekely_Suppl_Table_S3.xls: This file contains the outcome of the transcriptome-wide rhythmicity analyses on the kidney datasets (related to Figure 3A).

CasteloSzekely_Suppl_Table_S4.xls: This file contains the outcome of the rhythmicity analyses in kidney and liver for the 178 commonly rhythmic genes (related to Figure 3C).

Supp_dataset_S1.zip: This zip-file contains transcriptome-wide, time-resolved expression plots for kidney RPF (blue) and RNA (orange) in the left panels (with “error bars” connecting the two replicates of each timepoint) and TE in the right panels.

Supp_dataset_S2.zip: Similar to Supp_dataset_S2.zip, expression plots for kidney and liver for the 178 common rhythmic genes of Figure 3C.

### Author contributions

VCS and PJ performed the experiments. VCS and ABA analysed the data. DG and VCS wrote the manuscript with inputs from the others authors. DG designed the project.

## SUPPLEMENTARY FIGURE LEGENDS

**Figure S1.**
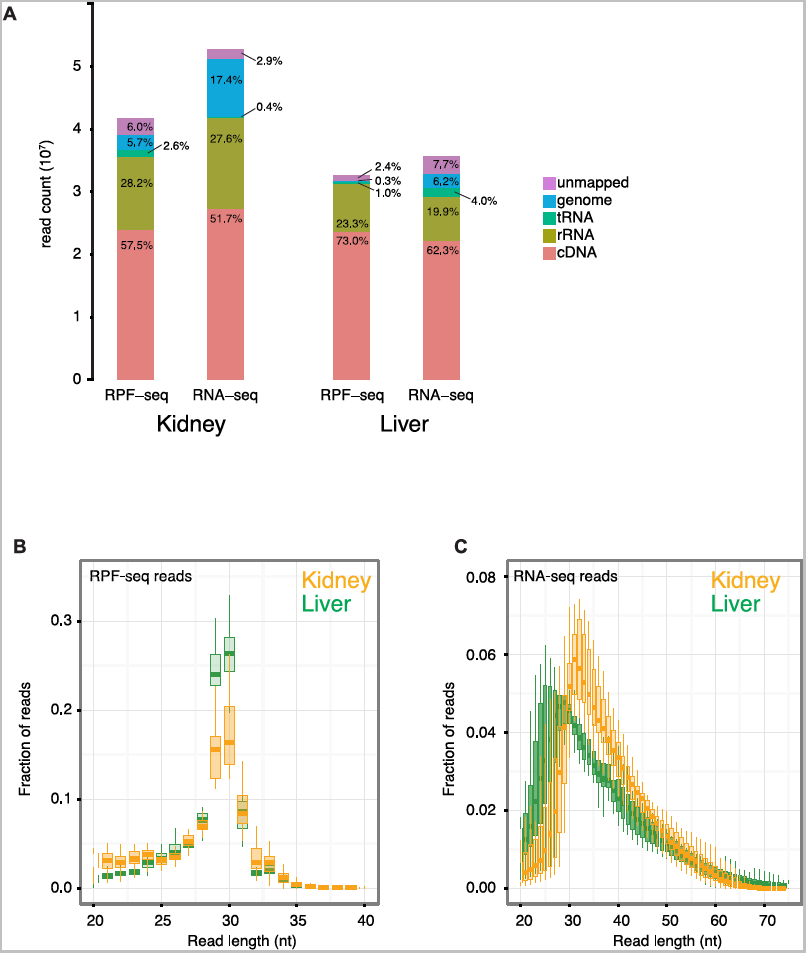
Overview of sequencing outcome and read length distribution for RPF- and RNA-seq data used in the study. **A**. Summary of outcome of the sequential mapping pipeline, indicating the number (y-axis) and percentage (within bars) of reads mapping to each database, averaged over all timepoints. For each sample of the four datasets an average of more than 20 million reads mapped to the protein-coding transcriptome (cDNA) and was used for the study. **B and C** RPF-seq (B) and RNA-seq (C) read length after trimming of adaptors showed that most RPF-seq reads had a length of 29-30 nucleotides in both organs, whereas RNA-seq fragments showed a broader distribution as expected from chemical RNA fragmentation. Boxplots represent the interquartile range and whiskers extend to the minimum and maximum values within 1.5 times the interquartile range.

**Figure S2.**
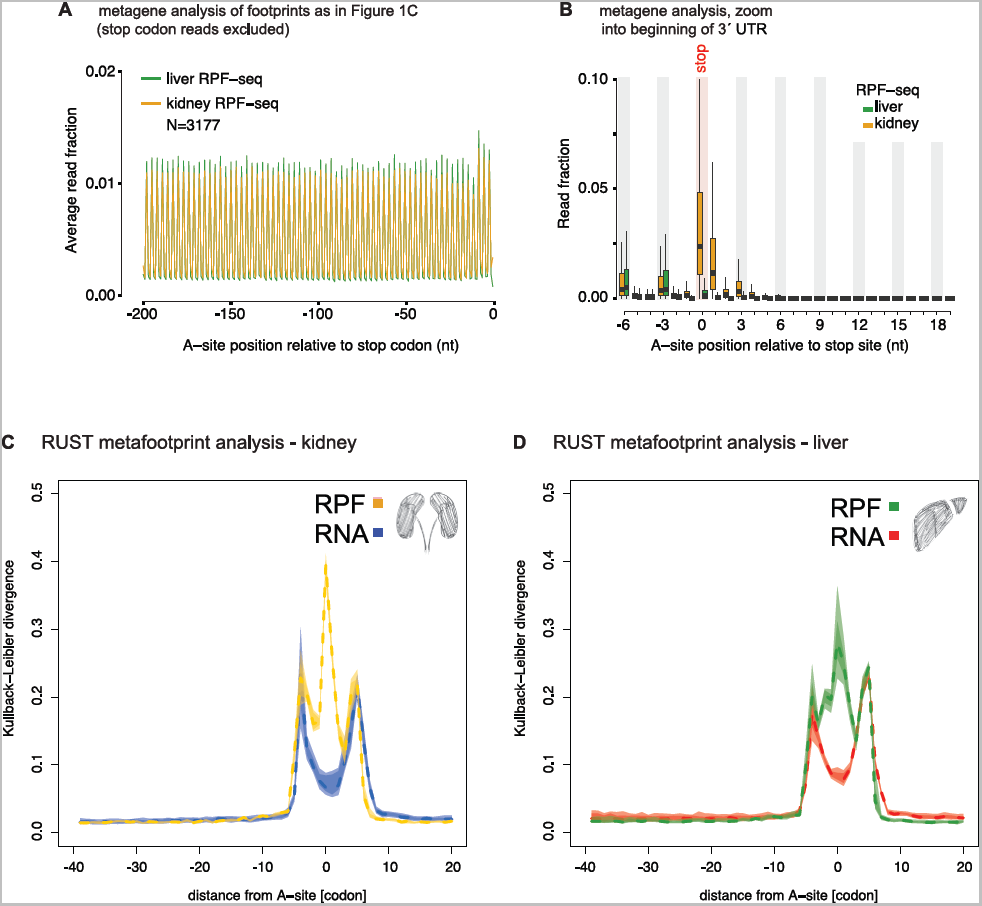
RPF reads at the stop codon and quality control. **A** A-site position of RPF-seq reads in the last 200 nt of the CDS, excluding stop codon reads. Read density along the CDS was similar in liver and kidney; thus the higher read density observed in Fig. 1C at the stop codon does not affect CDS-based calculations. **B** Zooming into the footprint read density at the end of the CDS, stop codon, and beginning of the 3′ UTR indicated read differences between organs for the stop codon itself and up to 4 nt downstream, which were increased in kidney. Stop codon reads are counted towards the 3′ UTR and their higher level in kidney thus also explains why our analysis in Fig. 1B shows more 3′ UTR reads for this organ. The remainder of the 3′ UTR shows a similar depletion of reads in both organs. **C and D** Ribo-seq Unit Step Transformation (RUST) metafootprint analyses toevaluate the contribution of local mRNA positions to the density of footprints. Light and dark coloured polygonal areas denote the 10%-90% and 25%-75% percentiles, respectively, and the dash-line denotes the median of the Kullback–Leibler divergence (K–L) profiles of all samples within kidney (C) and liver (D) separately for RPF and RNA reads (colour code in inset). The K-L profile for each sample was calculated from the RUST ratio values of 61 sense codons across a moving window of 40 triplet codons upstream to 20 triplet codons downstream of predicted A-sites. High K-L divergence maxima (lowest relative entropy or highest information gain) are thus found in the vicinity of the A-site in RPF libraries, and the 5′ and 3′ termini of the reads in RNA libraries. Importantly, the profiles are similar for kidney and liver, indicating overall similar footprint quality in the two independent datasets.

**Figure S3.**
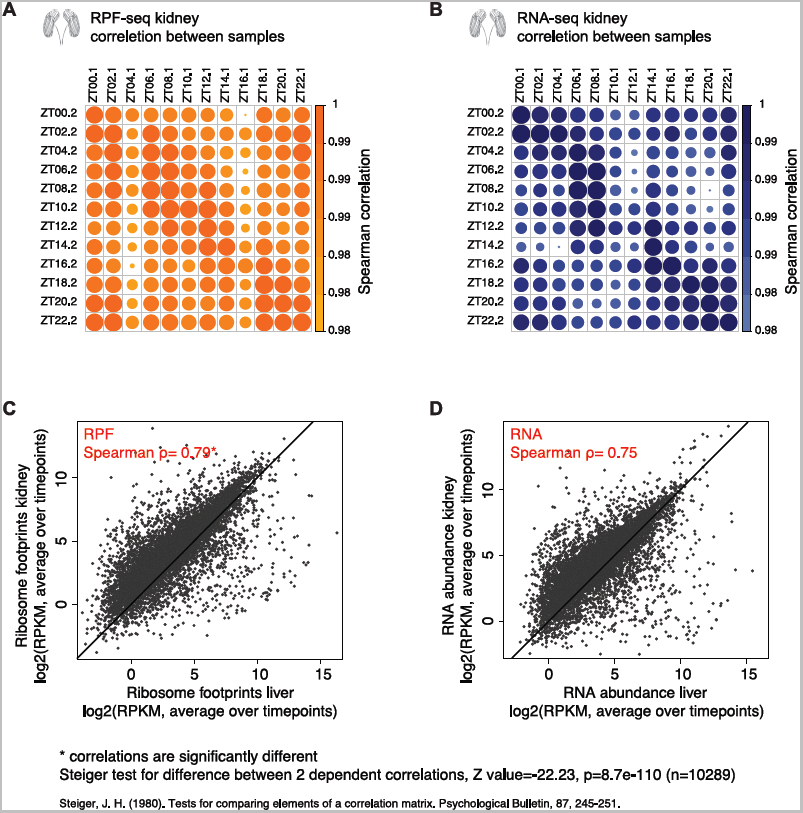
High technical and biological reproducibility of datasets. **A and B** Spearman correlation of normalised CDS read counts between timepoints and between replicates for kidney RPF-seq (A) and kidney RNA-seq (B) datasets. The correlation coefficient is indicated by the size and shading of the disks. Biological replicates thus show excellent correlation; moreover, the correlation coefficients of different timepoints reflect the rhythmic nature of the data. **C and D** Normalised CDS read counts (RPKM) in liver vs. kidney at the RNA (C) andRPF level (D). In these graphs, the averages over all timepoints were compared for the set of commonly expressed genes (N=10289; see Fig. 2A). Note the overall higher Spearman correlation at the footprint level than transcript level. The difference between correlations is highly significant with p=8.7e-110, Z value=-22.23; Steiger test for difference between 2 dependent correlations (Reference: Steiger JH, 1980. Psychological Bulletin, 87, 245-251).

**Figure S4.**
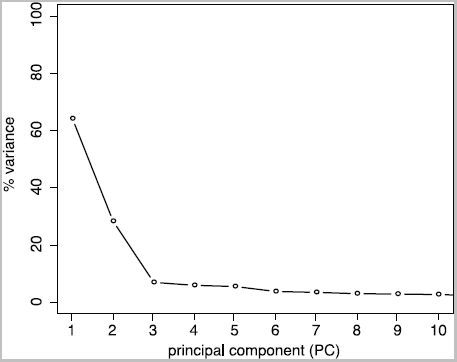
Additional information for Principal Component Analysis. Scree plot showing the first 10 components of the PCA in Fig. 1D-E. Components 1 and 2 explained most variance, followed by PC3 to PC5, which explained a closely similar proportion of variance in the data; the plateau was apparent from the sixthcomponent.

**Figure S5.**
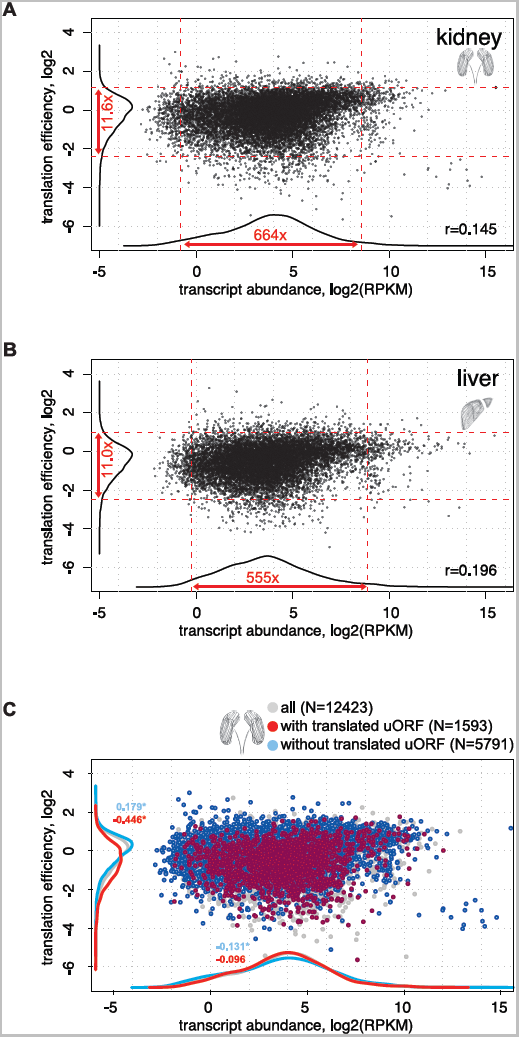
Contribution of translation efficiency to overall gene expressionvariation within organs. **A and B** Scatterplot of mRNA abundance vs. translation efficiency (TE) in kidney (A;N=12423 genes) and liver (B; N=10676 genes), averaged over all timepoints. Corresponding density lines are plotted on the margins. Dotted red lines represent the 2.5 and 97.5 percentiles, and the corresponding fold change is indicated. The transcript abundance range for 95% of genes thus spanned two orders of magnitude (>500-fold range in either organ), whereas TE dynamic range was less than 12-fold in either organ. Transcript abundance differences can thus be considered the main source of gene expression variability in the tissues. Moreover, Pearson’s r values of 0.145 (kidney) and 0.196 (liver) indicate weak positive correlation of transcript abundance and TE. **C** Scatterplot of transcript abundance (TA) vs. translation efficiency (TE) in mainCDS for kidney and as averages over all timepoints. Highlighted are single protein-coding genes that contain (red) or do not contain (blue) translated uORFs. Corresponding density lines are plotted on the margins. uORF translation is thus clearly associated with significantly reduced translation efficiency. Numbers on the density curves indicate the location shift relative to all transcripts. Genes with translated uORFs: TA, p=0.16; TE, p<2.2e-16 (Wilcoxon rank sum test). Genes without translated uORFs: TA, p=8.7e-5; TE, p<2.2e-16 (Wilcoxon rank sum test).

**Figure S6.**
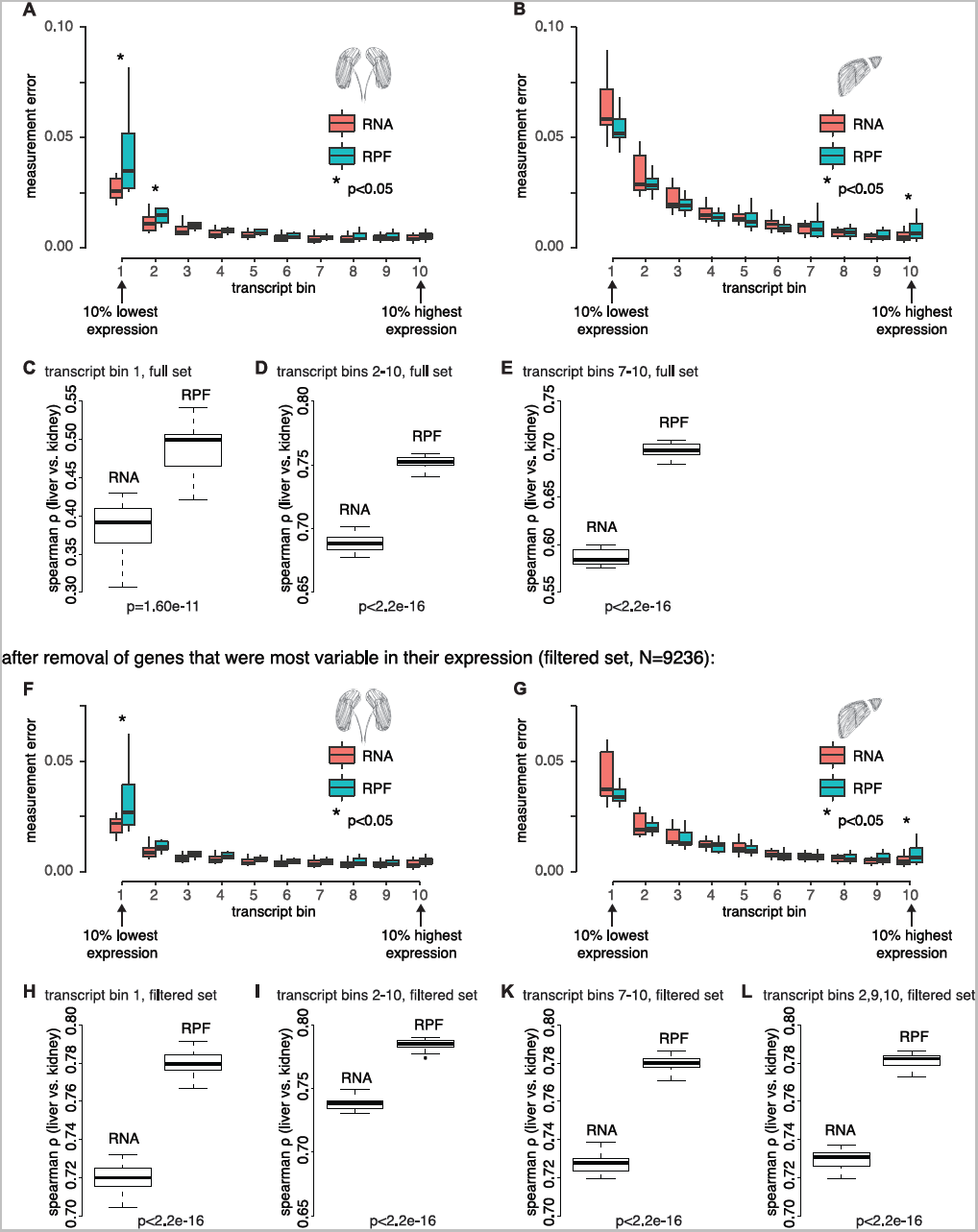
Translational compensation is independent of technical biases inthe datasets. **A and B** Measurement error (ME) of all genes in the dataset (n=10289) wascalculated separately for RNA-seq (red) and RPF-seq (turquoise) for kidney (A) and liver (B), and plotted as a function of increasing average expression levels. Briefly, measurement errors within each bin were calculated as in Albert et al., 2015 (Albert FW, Muzzey D, Weissman JS, Kruglyak L, 2014. PLoS Genetics, 10, e1004692) using the 12 timepoints as replicates for the measurement estimates (see Methods). **C to E** Spearman correlation between liver and kidney for RNA and RPF data for (C)genes showing the highest measurement errors (bin 1 in A, B), (D) bins 2-10, and (E) genes that have higher measurement error in RPF-seq than in RNA-seq samples in both organs (bins 7-10). Each boxplot contains the correlation coefficients between organs for each timepoint and replicate sample. Together these analyses showed that RPF-seq samples have a higher degree of similarity across organs (indicated p-values are from paired t-tests on Fisher-transformed correlation coefficients), even when considering lowly expressed genes with higher associated experimental error (C), or when considering genes with higher RPF-seq than RNA-seq measurement errors (E). These results thus ruled out that a systematic lower measurement error in RPF-seq experiments could have been the underlying cause of the higher correlation in RPF-seq than RNA-seq observed in Fig. 2C. **F and G** Same as A and B, but with a filtered gene set in which specifically thosegenes that showed very different expression levels/high variability between organs or between datasets (RPF-seq, RNA-seq) were removed (see Methods). The reason to also analyse such a filtered set was that we wished to be sure that genes that were widely different in their gene expression level were not distorting the analyses (e.g. specifically causing extreme measurement errors under a condition where expression was very low). Moreover, because the binning into the groups was based on expression level across all sets (calculated as the fourth root of the product of liver RNA-seq, liver RPF-seq, kidney RNA-seq and kidney RPF-seq), the highly variable genes made binning inaccurate. This filtered set thus contained genes with overall better comparability across datasets; of note, the distribution of ME differences using the filtered set was very similar to the full set in A-B. **H to L** Inter-organ Spearman correlation in RNA-seq and RPF-seq samples forvarious gene bins as indicated, using the filtered set. As in C-E, even when considering for example the genes with the highest overall measurement error (H), or the genes with higher RPF-seq than RNA-seq measurement errors in both organs (L), a significantly higher correlation is observed in RPF-seq samples (paired t-test on Fisher-transformed correlation coefficients).

**Figure S7.**
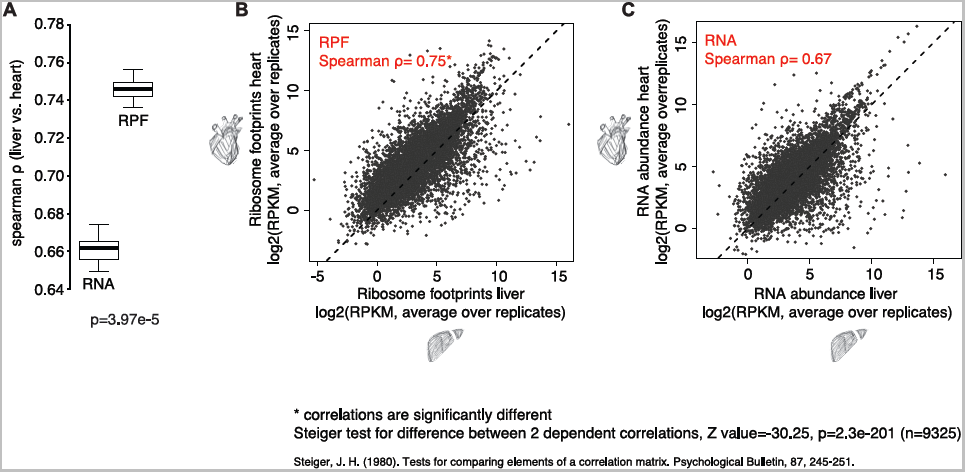
Translational compensation detected in rat liver and heart. **A)** Spearman correlation coefficient between rat heart and liver samples calculated from the data of Schafer et al., 2015 (Schafer S, Adami E, Heinig M, Rodrigues KE, Kreuchwig F, Silhavy J, van Heesch S, Simaite D, Rajewsky N, Cuppen E, Pravenec M, Vingron M, Cook SA, Hubner N, 2015. Nature Communications 8, 7200). Each boxplot contains the correlation coefficients of all possible pairwise comparisons between heart and liver replicates (remark: in this study, organs in each of the five replicates did not necessarily come from the same animals, thus precluding defined pairwise comparisons between same animals). The indicated p-value is the 95th percentile of the ensemble of p-values resulting from all possible comparisons between RPF-seq and RNA-seq correlation coefficients (paired t-test of Fisher-transformed coefficients). This analysis extended our observation of a globally higher conservation between organs at the level of translational output (protein production) than at the level of transcript abundance. **B and C** Normalised CDS read counts (RPKM) in rat liver vs. heart at the RPF-seq (B) and RNA-seq level (C), averaged over the five replicates used in the study of Schafer et al., 2015. Note the overall higher Spearman correlation at the footprint level as compared to the mRNA level. The difference between correlations is highly significant with p=2.3e-201, Z value=-30.25; Steiger test for difference between 2 dependent correlations (Reference: Steiger JH, 1980. Psychological Bulletin, 87, 245-251).

**Figure S8.**
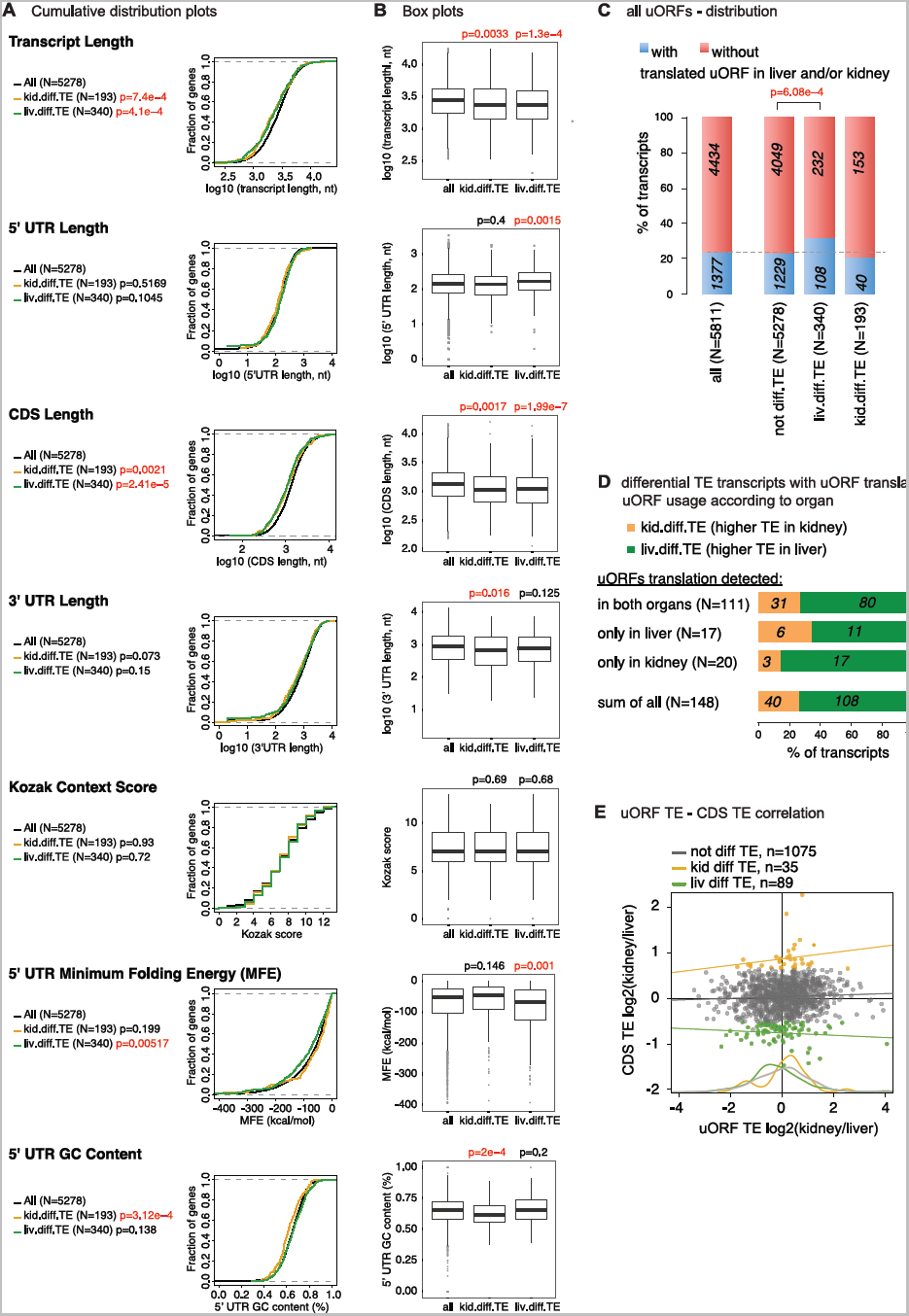
Analysis of transcript features with predictive value for differential TE. **A** Cumulative distribution of the indicated transcript features for single isoform genes that do not show differential TE (black, n= 5278), or that show differential TE and either higher TE in kidney (yellow, N=193) or in liver (green, N=340). The indicated p-values are Kolmogorov-Smirnov test results of each group vs. all. Statistically significant comparisons marked in red. **B** Same as (A), but in the form of boxplots and using Wilcoxon rank-sum test for the differences between group means (again, marked in red, significant results). **C** Fraction of single isoform genes with (blue) or without (red) translated uORFs in either organ. The group with differential and higher TE in liver contained significantly more translated uORF-containing transcripts than the genes not showing differential TE in either organ (p=6.08e-04; Fisher’s exact test); for kidney, there was a slight depletion of uORF-containing transcripts (non-significant). This analysis indicates that uORF usage may play a role in setting TE differences across tissues. Note that in this analysis, we did not yet distinguish whether the uORF was translated in liver and/or kidney, but we treated the 1377 genes with a translated uORF in at least one organ as a single group. **D** Organ-specific uORF usage and its association with differential TE. The group of genes with uORFs specifically translated in liver was enriched for transcripts better translated in in kidney, and vice versa, consistent with a role of tissue-specific uORF usage in setting TE differences. However, due to the low number of differential TE genes exhibiting uORF translation that was exclusive to one organ for this analysis, the enrichments and depletions did not reach statistical significance. **E** Scatterplot of upstream ORF vs. CDS TE differences across organs for genes containing translated uORFs in both organs and detected as differential TE with higher TE in kidney (yellow) or liver (green), or not showing differential TE (grey). An anticorrelation between uORF usage and CDS TE was only observed for genes with differential and higher TE in liver.

**Figure S9.**
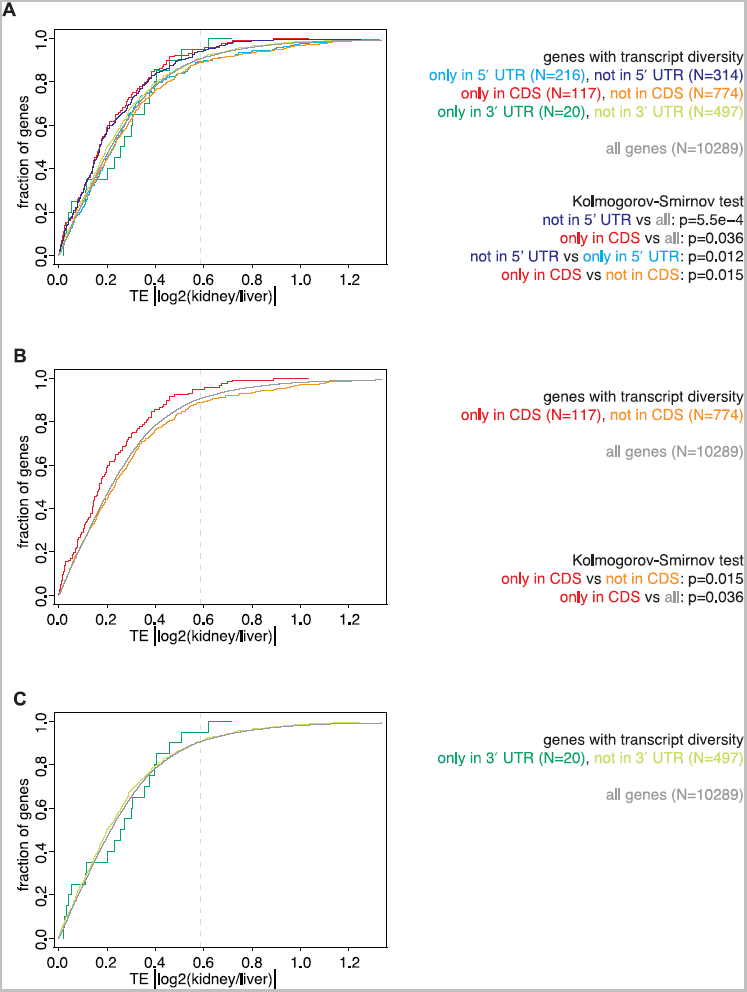
Relationship between transcript diversity and differential TE. **A** Cumulative distribution of the absolute kidney-to-liver TE ratio for genes whose transcript diversity is present or absent only in the indicated feature. The vertical dotted grey line marks the 1.5-fold difference used to define differential TE. In this Figure, all 7 groups are plotted (transcript diversity only/not in 5′ UTR, CDS, 3′ UTR; all genes); for better visibility, the analyses of individual features are also shown in separate panels, i.e. in Fig. 2G (5′ UTR), Fig. S9B (CDS) and Fig. S9C (3′ UTR). Collectively, these results showed that transcript diversity that originated only within the CDS (red), or that was excluded from the 5′ UTR (purple), or that was present only within the 3′ UTR (dark green), all showed smaller TE differences across organs, thus pointing towards variability within the 5′ UTR as a contributor to tissue-specific TE. **B** As in (A) showing the genes with transcript diversity present (red) or absent (orange) only in the CDS. Note that when transcript diversity is only in CDS (i.e. UTRs are identical), there is a significant shift to more similar TEs in both organs. This is consistent with the specific association of 5′ UTR diversity with differential TE that is shown in Fig. 2G. **C** As in (A) showing genes with transcript diversity present (dark green) or absent (light green) only in the 3′ UTR.

**Figure S10.**
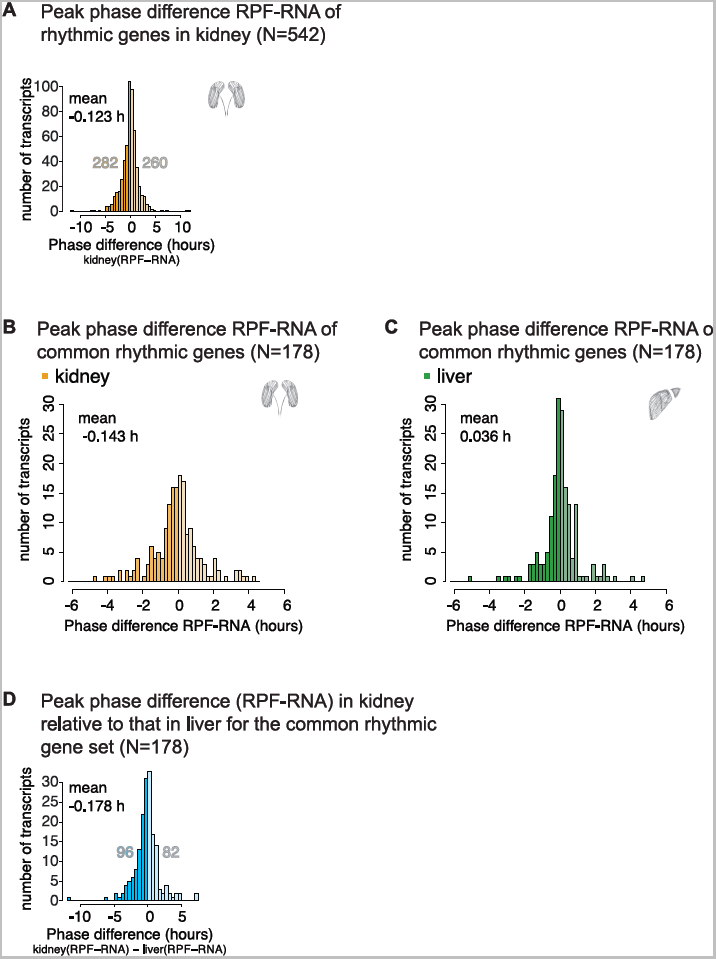
Analysis of phase differences in RNA and RPF rhythms in kidney and across organs. **A** Histogram of phase differences (RPF – RNA, in hours) for all genes that were detected as rhythmic in the kidney RPF and RNA data (N=542; see Fig. 3A). Although the distribution mean was not significantly different from 0, more genes had their footprint abundance peak advanced (N=282) than delayed (N=260) with respect to their mRNA abundance peak. **B and C** Histogram of the phase differences (footprints to mRNA abundance, in hours) in kidney (B) and liver (C) for the 178 genes rhythmic in both organs (gene set shown in Fig. 3C). We observed a broader distribution of phase differences in kidney and globally a phase advance of RPF with respect to RNA (-0.143 hours), as compared to overall stronger phase coherence of RPF and RNA in liver. See also Figures 3D-E. **D** Histogram of the differential (kidney – liver) phase difference (RPF – RNA) for the178 genes that were rhythmic throughout (Fig. 3C). Although statistically not reaching significance, the mean of -0.178 hours and the overall more genes for which the phase difference had negative values (96 vs. 82 genes) were consistent with the finding that RPF rhythms peak earlier than RNA rhythms specifically in kidney.

**Figure S11.**
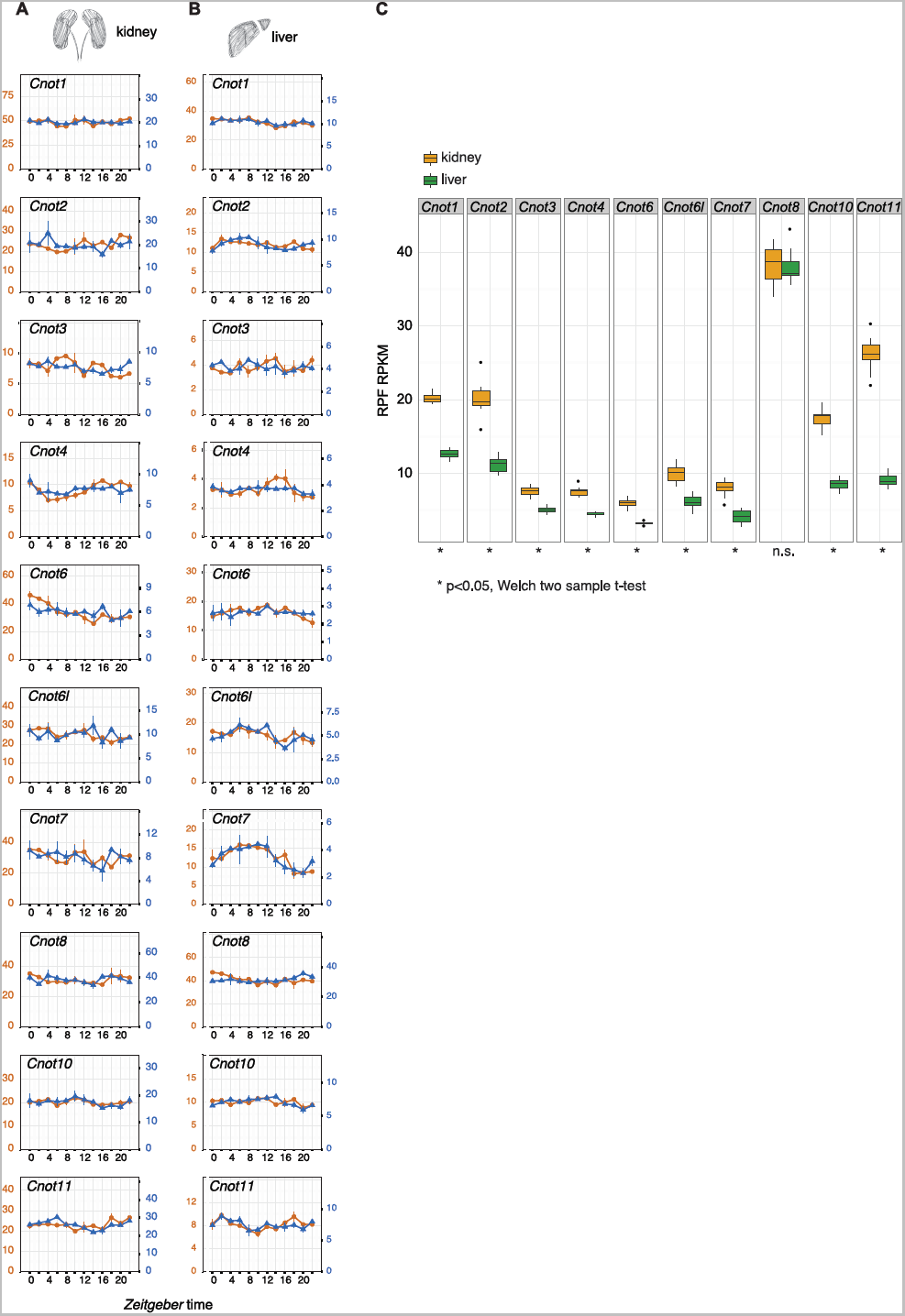
Higher expression of deadenylase complex subunits in kidney. **A and B** Daily expression profiles of the CCR4-NOT complex components in kidney (A) and liver (B) at the RNA (orange) and RPF (blue) level. **C** RPF expression of the CCR4-NOT subunits (averages over the day). Boxplots represent the interquartile range and whiskers extend to the minimum and maximum expression within 1.5 times the interquartile range. Note that differences in protein biosynthesis are statistically significant for all subunits (p<0.05, two-sample t-test) apart from *Cnot8*.

**Figure S12.**
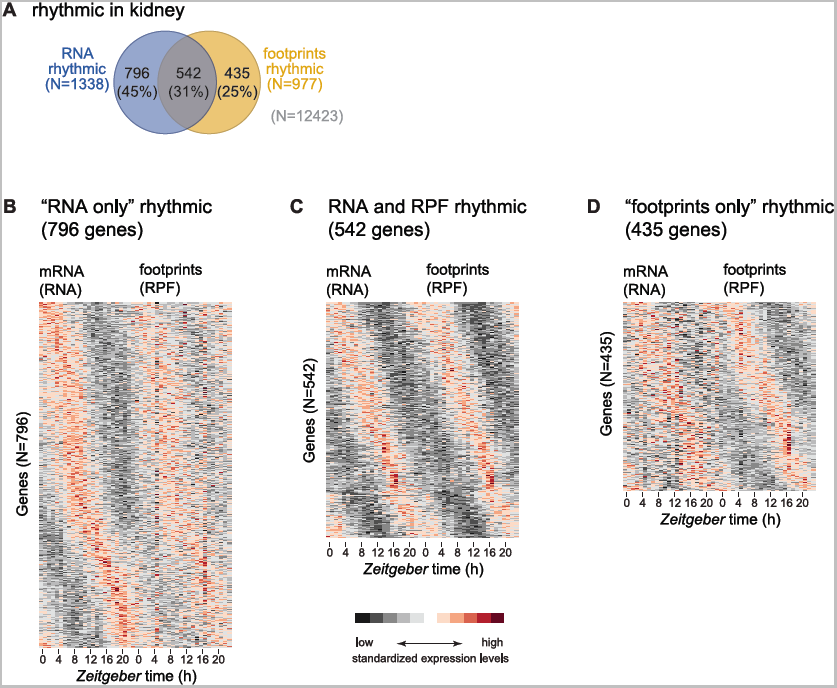
Heatmaps of all detected RPF and RNA rhythms indicate false-negatives of the rhythmicity detection method. **A** Same as Fig. 3A, but re-plotted here for ease of comparison with (B-D). **B-D** Heatmap of RNA-seq (left) and RPF-seq (right) expression for genes detected as rhythmic only at the mRNA level (B, N=796), at both levels (C, N=542), and at the ribosome footprints only (D, N=435) in kidney. Gene expression levels are standardised by row (gene). Please note that even the panels that should represent “non-rhythmicity” (i.e. right panel in B and left panel in D) clearly showed underlying rhythmicity, albeit with more noise and/or lower amplitude. Many of these cases were therefore probably not truly “non-rhythmic” but rather false-negatives of the detection method (see Results section).

**Figure S13.**
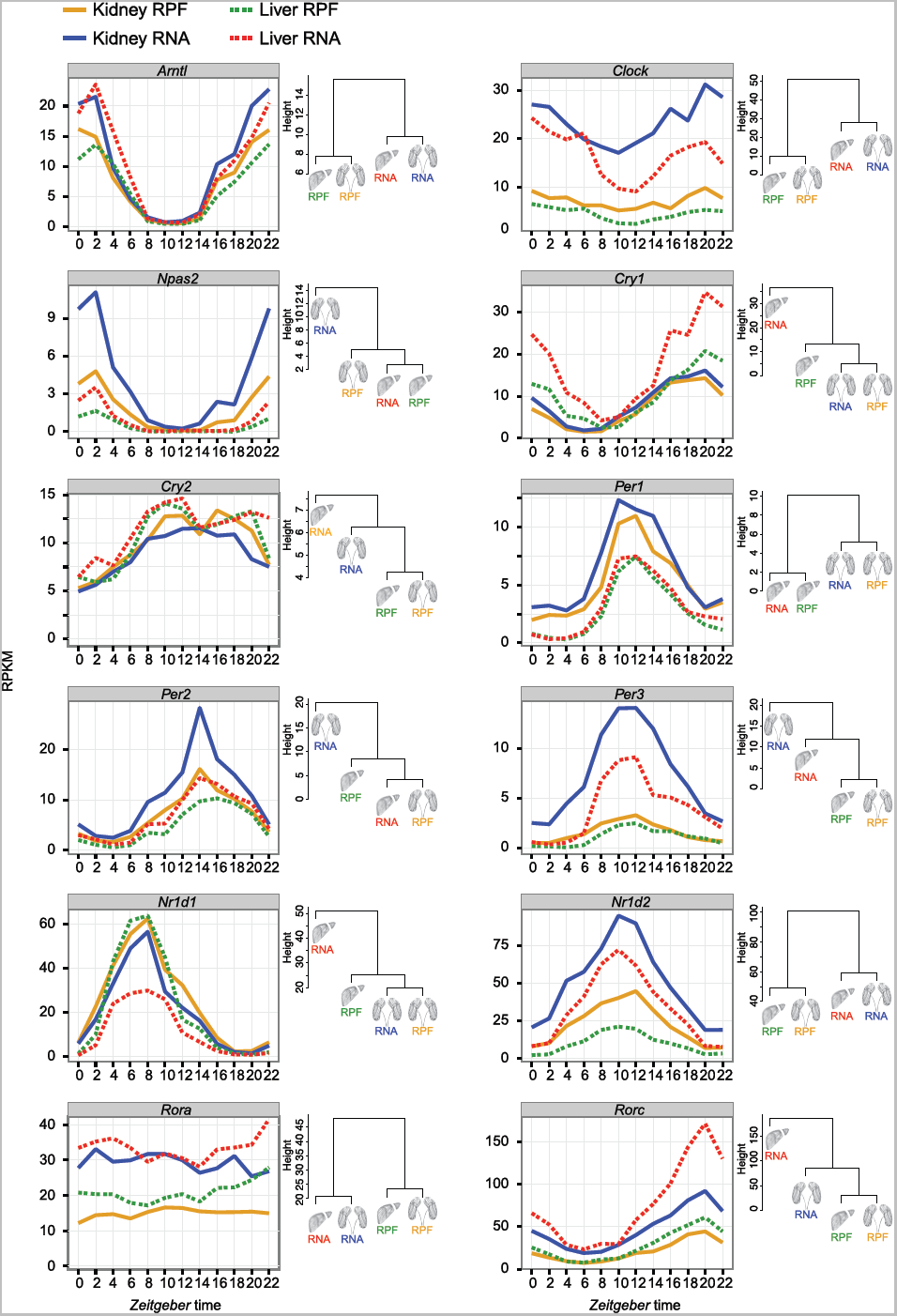
Core clock gene expression at RNA and RPF levels in both organs. **Left panels:** Daily expression profiles of the 12 main core clock genes shown in Fig. 5A-C. **Right panels:** Hierarchical clustering of the organs’ RNA and RPF profiles for each clock gene. Branch height represent the average Euclidean distance. Note that for 7 out of the 12 core clock genes, protein synthesis profiles were more conserved across organs than mRNA abundance and than RPF-RNA within organs.

**Figure S14.**
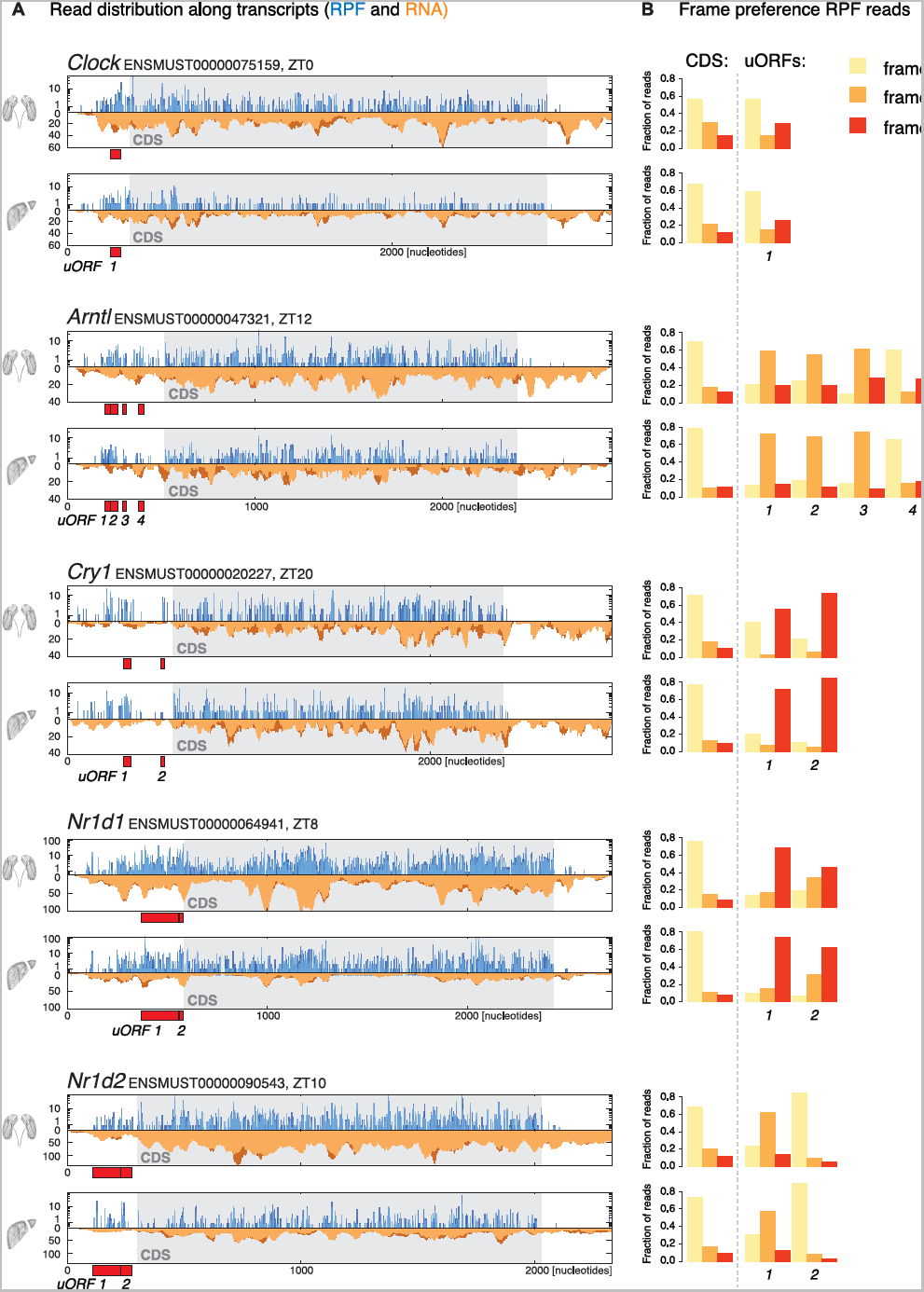
Read distribution for uORF-containing core clock genes. **A** Normalised read distribution for RPF (in blue) and RNA (in orange) along core clock transcripts containing uORFs in kidney (top) and liver (bottom) for the timepoint of maximal CDS translation. Red boxes indicate AUG-initiated uORFs as predicted in our analyses. For scaling issues and better visualisation, only a portion of the 3′ UTRs, corresponding to the same length as the full 5′ UTR, is depicted (exception *Nr1d1*, for which the 3′UTR is so short that it is shown full length). **B** Read distribution to the three translation frames showed a frame bias of footprint reads for most predicted uORFs that was in a similar range as the frame bias on the CDS. This frame preference is indicative of active translation on the uORFs.

